# Interferon lambda restricts herpes simplex virus skin disease by suppressing neutrophil-mediated pathology

**DOI:** 10.1101/2023.09.11.557277

**Authors:** Drake T. Philip, Nigel M. Goins, Nicholas J. Catanzaro, Ichiro Misumi, Jason K. Whitmire, Hannah M. Atkins, Helen M. Lazear

## Abstract

Type III interferons (IFN-λ) are antiviral and immunomodulatory cytokines that have been best characterized in respiratory and gastrointestinal infections, but the effects of IFN-λ against skin infections have not been extensively investigated. We sought to define the skin-specific effects of IFN-λ against the highly prevalent human pathogen herpes simplex virus (HSV). We infected mice lacking the IFN-λ receptor (*Ifnlr1*^-/-^), both the IFN-λ and the IFN-αβ receptor (*Ifnar1*^-/-^ *Ifnlr1*^-/-^), or IFN-λ cytokines (*Ifnl2/3*^-/-^) and found that IFN-λ restricts the severity of HSV-1 and HSV-2 skin lesions, independent of a direct effect on viral load. Using conditional knockout mice, we found that IFN-λ signaling in both keratinocytes and neutrophils was necessary to control HSV-1 skin lesion severity, and that IFN-λ signaling in keratinocytes suppressed CXCL9-mediated neutrophil recruitment to the skin. Furthermore, depleting neutrophils or blocking CXCL9 protected against severe HSV-1 skin lesions in *Ifnlr1*^-/-^ mice. Altogether, our results suggest that IFN-λ plays an immunomodulatory role in the skin that restricts neutrophil-mediated pathology during HSV infection, and suggest potential applications for IFN-λ in treating viral skin infections.

**Graphical Abstract:** 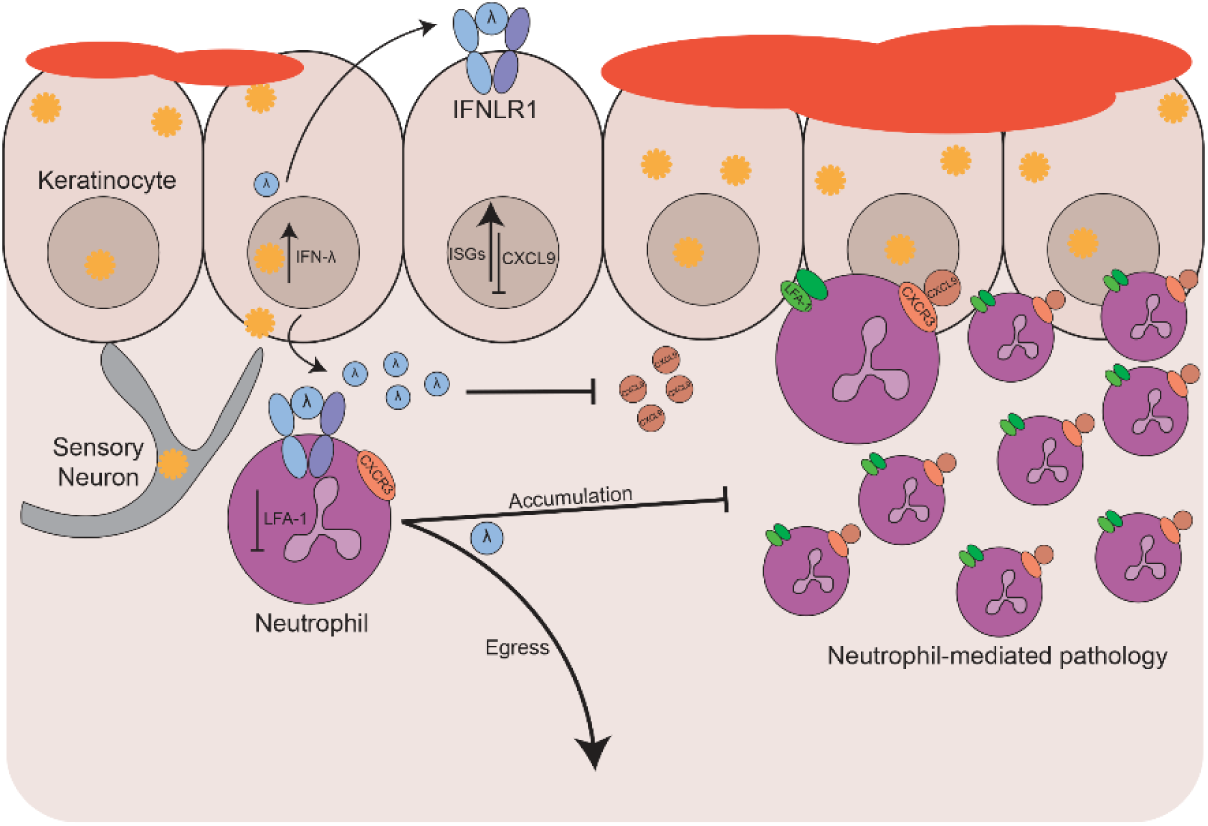

## INTRODUCTION

The skin is one of the largest physiologic and immunologic barriers that separates the host from its environment and continuously encounters commensal and pathogenic microbes. Dysregulation of this barrier can result in inflammatory skin conditions, such as atopic dermatitis (eczema), or susceptibility to viral, bacterial, and fungal infections (Handfield et al., 2018). Resident and infiltrating immune cells as well as barrier-specific cytokines contribute to providing a balanced immune response in the skin (Nestle et al., 2009). The skin is comprised of two layers: the epidermis and dermis. The epidermis primarily consists of keratinocytes (skin epithelial cells) and T cells (CD8 and γδ). Keratinocytes form the physical barrier of the skin but also are immunologically active, coordinating resident and infiltrating leukocyte responses (Nestle et al., 2009). The dermis is primarily made of fibroblasts and is surveyed by a more diverse set of immune cells including macrophages, dendritic cells, granulocytes, Langerhans cells, and T cells (CD8, CD4, γδ) (Nestle et al., 2009). Both layers of the skin act in concert to mediate immune responses, using a variety of cytokines and chemokines—including interferons (IFNs)—in balancing pathogen clearance with homeostasis and barrier integrity (Kobayashi et al., 2019).

There are three families of IFNs: type I (IFN-α/β), type II (IFN-γ), and type III (IFN-λ), which are distinguished by their receptor usage (Mesev et al., 2019). IFN-α/β and IFN-γ typically mediate systemic antiviral and immunomodulatory functions whereas the activity of IFN-λ is most evident at epithelial barriers such as the respiratory and gastrointestinal tracts (Lazear et al., 2019). There are four IFN-λ subtypes in humans (IFN-λ1-4), whereas mice express only IFN-λ2 and IFN-λ3 (Lazear et al., 2019). All IFN-λ subtypes signal through the same heterodimeric receptor comprised the IFNLR1 and IL10Rβ subunits (Dowling & Forero, 2022). IFN-λ intracellular signaling is similar to IFN-α/β in that upon receptor binding Jak1 and Tyk2 are activated to phosphorylate STAT1 and STAT2, which bind to IRF9 and translocate to the nucleus to drive interferon stimulated gene (ISG) expression (Dowling & Forero, 2022). Although IFN-λ and IFN-α/β canonically share the same intracellular signaling cascade and activate similar antiviral transcriptional programs, there has been a growing appreciation for their different antiviral and immunomodulatory functions. Their difference in function is in part due to kinetics, as IFN-α/β is induced more rapidly upon infection whereas IFN-λ is produced later and for longer throughout infection (Lazear et al., 2019; Mesev et al., 2019). Further, whereas the IFN-α/β receptor (IFNAR1/IFNAR2 heterodimer) is expressed ubiquitously, IFNLR1 expression is restricted primarily to epithelial cells and select leukocytes such as neutrophils, dendritic cells, and macrophages (Lazear et al., 2019). The high expression of IFNLR in epithelial tissues has led to substantial characterization of IFN-λ antiviral and immunomodulatory functions at epithelial barriers including the respiratory tract and gastrointestinal tract. However, relatively little is known about the role of IFN-λ in the skin, even though the skin is an important epithelial barrier and home to a variety of IFN-λ producing and responding cell types.

Therefore, we sought to investigate the functions of IFN-λ signaling in the skin using a highly prevalent human skin pathogen, herpes simplex virus (HSV-1 and HSV-2). HSV-1 primarily manifests as orofacial or genital lesions but severe complications of HSV-1 infection include herpes encephalitis, herpes keratitis, and eczema herpeticum (Kollias et al., 2015). HSV-2 most commonly causes genital herpes and can cause severe disease when transmitted to neonates at delivery (Bradley et al., 2014). HSV-1 and HSV-2 establish life-long persistent (latent) infections in sensory neurons that are never cleared by the immune system; viral reactivation from latency results in recurrent epithelial lesions (cold sores, genital herpes) and transmission to new individuals. Over half of the US adult population are seropositive for HSV-1 and 15% are seropositive for HSV-2 (Bradley et al., 2014). The disease associated with HSV-1 infection, in both mice and humans, is partly immune-mediated (Kollias et al., 2015) but the viral and host factors that drive disease severity are not fully understood, including the effects of IFN-λ signaling in HSV-1 skin infection.

Here, we report a protective role for IFN-λ signaling in restricting severe HSV-1 skin disease. Using a mouse model of HSV-1 skin infection, we found that IFN-λ-dependent protection from severe HSV-1 skin disease requires signaling in both keratinocytes and leukocytes (including neutrophils). We found that neutrophils are the primary leukocytes recruited to the skin during HSV-1 infection, and that mice lacking IFN-λ signaling (*Ifnlr1^-/-^*) have more skin-infiltrating neutrophils compared to wild-type mice. We found a greater proportion of neutrophils expressing the tissue homing integrin leukocyte function associated antigen-1 (LFA-1) in *Ifnlr1^-/-^* compared to wild-type mice, suggesting that *Ifnlr1^-/-^* neutrophils are highly recruited to and retained in HSV-1 infected skin. Furthermore, we showed that depleting neutrophils prevented the development of severe HSV-1 skin lesions in *Ifnlr1*^-/-^ mice and that IFN-λ signaling in keratinocytes limits CXCL9-mediated neutrophil recruitment to the skin. Altogether, our results suggest that IFN-λ plays an immunomodulatory role in the skin that restricts neutrophil-mediated pathology during HSV infection and suggest potential applications for IFN-λ in treating viral skin infections.

## RESULTS

### Defining HSV-1 skin lesion severity in the mouse flank infection model

Flank scarification is a well-established mouse model for studying HSV-1 pathogenesis (Simmons & Nash, 1984; van Lint et al., 2004; Wang et al., 2019). In the standard model, HSV-1 is scratched into depilated flank skin using a 27 gauge needle (Simmons & Nash, 1984; van Lint et al., 2004; Wang et al., 2019). The virus replicates at the inoculation site, infects innervating sensory neurons and traffics in a retrograde direction to the neuron cell body located in the dorsal root ganglia (DRG), then traffics in an anterograde direction, returning to the entire dermatome innervated by that DRG, producing a dermatome skin lesion (Fig. S1A) (Wang et al., 2019). To improve the reproducibility of this infection model, we tested two modifications to the commonly used procedures. First, we depilated mice by manual plucking, rather than shaving plus chemical depilation (e.g. Nair cream). Plucking re-sets the hair cycle (Kobayashi et al., 2019), producing an immunologically uniform environment across the depilated skin, and also is faster and easier than shaving plus chemical depilation. We found no significant difference in dermatome lesion area for mice depilated by plucking compared to shaving plus Nair (Fig. S1B), suggesting that the method of depilation does not impact HSV-1 skin infection in this model, so further experiments depilated mice by plucking. Next, we compared three methods for disrupting the skin barrier, as HSV-1 does not efficiently infect intact skin. We applied 10^6^ focus-forming units (FFU) of HSV-1 (strain NS) onto depilated flank skin and abraded the inoculation site skin by i) 40 gentle scratches with a 27G needle, ii) 10 closely-spaced punctures with a Quintip skin test (allergy needle), or iii) a single puncture with a Greer pick skin test, and measured viral loads in dermatome skin at 6 days post-infection (dpi). We found that skin viral loads corresponded to the extent of skin barrier disruption, with needle scarification producing a higher median viral load than a Quintip puncture (5.0 vs. 4.2 Log_10_ FFU/g) and no virus detected after a Greer pick puncture (Fig. S1C). Although needle scarification produced higher median viral loads, there was greater mouse-to-mouse variability compared to Quintip puncture (standard deviation (SD) 1.1 vs 0.7 Log_10_ FFU/g) and we and others have observed substantial operator-to-operator variability with needle scarification. Further, we found no difference in median lesion area between needle scarification and Quintip puncture (Fig. S1D); therefore, we used Quintip puncture to disrupt the skin barrier for further experiments.

We also sought to define a categorical system for classifying dermatome lesions as mild, moderate, or severe to better describe skin lesion phenotypes. We infected 20 wild-type mice with HSV-1, measured dermatome lesion area at 6 dpi, and found a mean area of 11 mm^2^ (SD 12 mm^2^) (Fig. S1E). We defined severe lesions as those with area >1 SD above the mean (>23 mm^2^), mild lesions as those with area smaller than the average size of the inoculation site (<5 mm^2^), and moderate lesions as 5-23 mm^2^ (Fig. S1F-G). We used both lesion area and categorical severity score to assess dermatome lesions in further experiments. Altogether, the flank infection model provides a robust system for assessing HSV-1 pathogenesis in the skin.

### IFN-λ signaling protects mice from severe HSV skin disease

IFN-λ provides antiviral protection at epithelial barriers such as the respiratory and gastrointestinal tracts (Broggi et al., 2020), but the effects of IFN-λ on viral infection in the skin have not been extensively investigated. To address this gap in the field, we infected wild-type mice and mice that lack the IFN-λ receptor (*Ifnlr1^-/-^*) with 10^6^ FFU of HSV-1 and evaluated skin lesion area, severity, and viral load 2-10 dpi (Fig. 1A-D). Experimental mice were generated by crossing mice with a floxed allele of the IFN-λ receptor (*Ifnlr1*^f/f^) with mice hemizygous for Cre recombinase expression under a ubiquitous promoter (*ActB*-Cre), producing littermate Cre-(wild-type) and Cre+ (*Ifnlr1^-/-^)* mice. Genotyping was performed retrospectively after data analysis and experiments were conducted blinded to the *Ifnlr1* status of each mouse. We found that *Ifnlr1^-/-^* mice developed significantly larger skin lesions than wild-type mice at 6 dpi, the time of peak disease (median lesion area 36.2 vs 7.7 mm^2^, *P* <0.01; Fig. 1B), and that a greater proportion of *Ifnlr1^-/-^* mice developed severe skin lesions compared to wild-type mice (71% vs 23%, *P* < 0.05; Fig. 1C). There was no significant difference in lesion area between *Ifnlr1^-/-^* and wild-type mice at 8 or 10 dpi (Fig. 1B). We measured viral loads by focus-forming assay from *Ifnlr1^-/-^* and wild-type skin lesions and from infected skin prior to lesion development (Fig. 1D). We found no significant differences in skin viral load between *Ifnlr1^-/-^* and wild-type mice at 2, 4, or 6 dpi (Fig. 1D), suggesting that IFN-λ signaling does not restrict HSV-1 replication in the skin. *Ifnlr1^-/-^* mice sustained detectable viral loads in the skin longer than wild-type mice (8 of 12 *Ifnlr1^-/-^* mice positive at 8 dpi vs 2 of 12 wild-type mice, *P* < 0.05), but *Ifnlr1^-/-^* and wild-type mice both cleared the infection from the skin by 10 dpi (Fig. 1D). To determine whether differences in dermatome skin lesions resulted from differences in sensory neuron infection, we infected wild-type and *Ifnlr1^-/-^* mice, harvested DRG at 6 dpi, and quantified viral loads by qPCR. We found no significant differences in HSV-1 viral genomes in DRG 6 dpi (Fig. 1E). Altogether, these results suggest that IFN-λ restricts HSV-1 skin lesion severity independently of a direct effect on viral replication.

**Figure 1.**
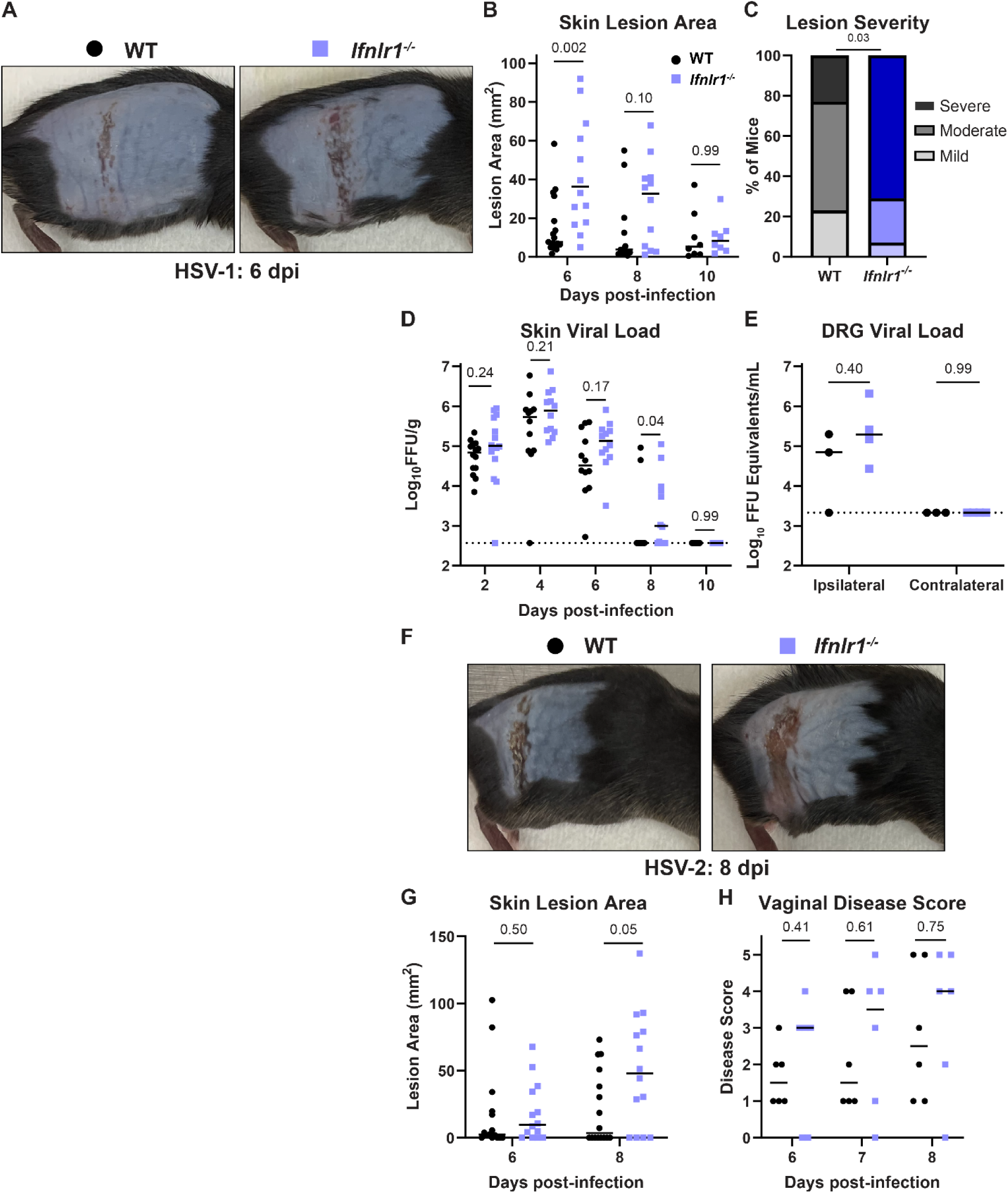
IFN-λ signaling protects against severe HSV skin disease without affecting viral replication. **A-E.** 8-12 week-old male and female WT and *Ifnlr1^-/-^* mice were infected with 10^6^ FFU of HSV-1 strain NS. **A**. Dermatome skin lesions were photographed 6 dpi. **B.** HSV-1 skin lesions were measured from photographs 6, 8, and 10 dpi using imageJ. **C.** Skin lesion severity was categorized based on 6 dpi lesion area. **D.** Skin viral loads were measured 2-10 dpi by focus-forming assay. **E.** Viral loads in ipsilateral and contralateral dorsal root ganglia (L2-L5 pooled for each mouse) were measured by qPCR. **F-G.** 8-12 week-old male and female WT and *Ifnlr1^-/-^* mice were infected with 10^3^ FFU of HSV-2 strain 333. **F.** Dermatome lesions photographed 8 dpi. **G.** HSV-2 skin lesions were measured from photographs 6 and 8 dpi using ImageJ. **H.** 8-12 week-old female WT and *Ifnlr1^-/-^* mice were infected with 10^3^ FFU of HSV-2 strain 333 intravaginally and HSV-2 vaginal disease was scored 6-8dpi as follows: 1: mild redness and swelling, 2: visible ulceration and fur loss, 3: severe ulceration and signs of sickness, 4: hindlimb paralysis, 5: moribund/dead. Differences in lesion area and viral load were compared by Mann-Whitney U test. Differences in categorical skin disease were compared by Cochran-Armitage test. *P* values are reported with *P* < 0.05 considered to be statistically significant.

We next tested whether IFN-λ had a similar protective effect against skin lesions caused by HSV-2. We infected mice with 1000 FFU of HSV-2 (strain 333) and measured dermatome lesion areas 6 and 8 dpi (Fig. 1F-G). The kinetics of HSV-2 infection were slower, with peak skin lesion area at 8 dpi, compared to 6 dpi for HSV-1, even though HSV-2 (333) was more virulent than HSV-1 (NS) (∼75% mortality at an inoculation dose of 1000 FFU of HSV-2 (333) compared to ∼0% mortality at an inoculation dose of 10^6^ FFU of HSV-1 (NS). Similar to HSV-1, *Ifnlr1*^-/-^ mice developed significantly larger lesions compared to wild-type mice (median lesion area at 8 dpi 47.8 mm^2^ vs 3.6 mm^2^, *P* < 0.05; Fig. 1G). Despite the slower kinetics of HSV-2 skin disease, the peak area of HSV-2 lesions in *Ifnlr1*^-/-^ mice was larger than HSV-1 lesions (median lesion area 47.8 mm^2^ at 8 dpi vs 36.2 mm^2^ at 6 dpi; Fig. 1B and 1G). We next asked whether IFN-λ protected against HSV disease at other epithelial sites. Although both HSV-1 and HSV-2 infect the genital tract in humans and mice, HSV-1 does not cause overt vaginal disease in mice. Therefore, we used HSV-2 to assess the effects of IFN-λ during vaginal infection. We found no significant difference between *Ifnlr1*^-/-^ and wild-type vaginal disease severity 6-8 dpi (Fig. 1H). After 8 dpi, mice either began to recover or were euthanized due to neurologic disease signs (hindlimb paralysis and hunching). Altogether these results show that IFN-λ protects against severe skin disease caused by both HSV-1 and HSV-2.

### The protective effect of IFN-λ in the skin does not require IFN-α/β signaling

IFN-λ and IFN-αβ signal through distinct receptors but canonically activate an overlapping JAK/STAT signaling cascade and transcriptional response (Dowling & Forero, 2022). To determine whether the protective effects of IFN-λ in the skin acted via cross-talk with IFN-α/β signaling, we infected mice lacking the IFN-αβ receptor (*Ifnar1^-/-^*) or both the IFN-αβ and IFN-λ receptors (*Ifnar1^-/-^ Ifnlr1^-/-^* double-knockout, DKO) with HSV-1 and evaluated survival, skin lesion area, severity, and viral loads (Fig. 2A-E). As expected, in the absence of IFN-αβ signaling, the kinetics of HSV-1 infection were faster and disease was more severe, with all mice succumbing 5-8 dpi (Fig. 2A). At 4 dpi, the first day mice developed lesions, *Ifnar1^-/-^ Ifnlr1^-/-^* DKO mice developed significantly larger skin lesions compared to *Ifnar1*^-/-^ mice (median lesion area 67.8 mm^2^ vs 19.7 mm^2^, *P* < 0.001; Fig. 2B-C) and a significantly larger proportion developed severe lesions (100% vs. 44%, *P* < 0.001; Fig. 2D). We found no significant difference in skin viral loads at 2 dpi or 4 dpi (Fig. 2E). Altogether, these data indicate that IFN-λ limits severe HSV-1 skin disease independently of IFN-α/β and independent of a direct antiviral effect on HSV-1 replication in the skin.

**Figure 2.**
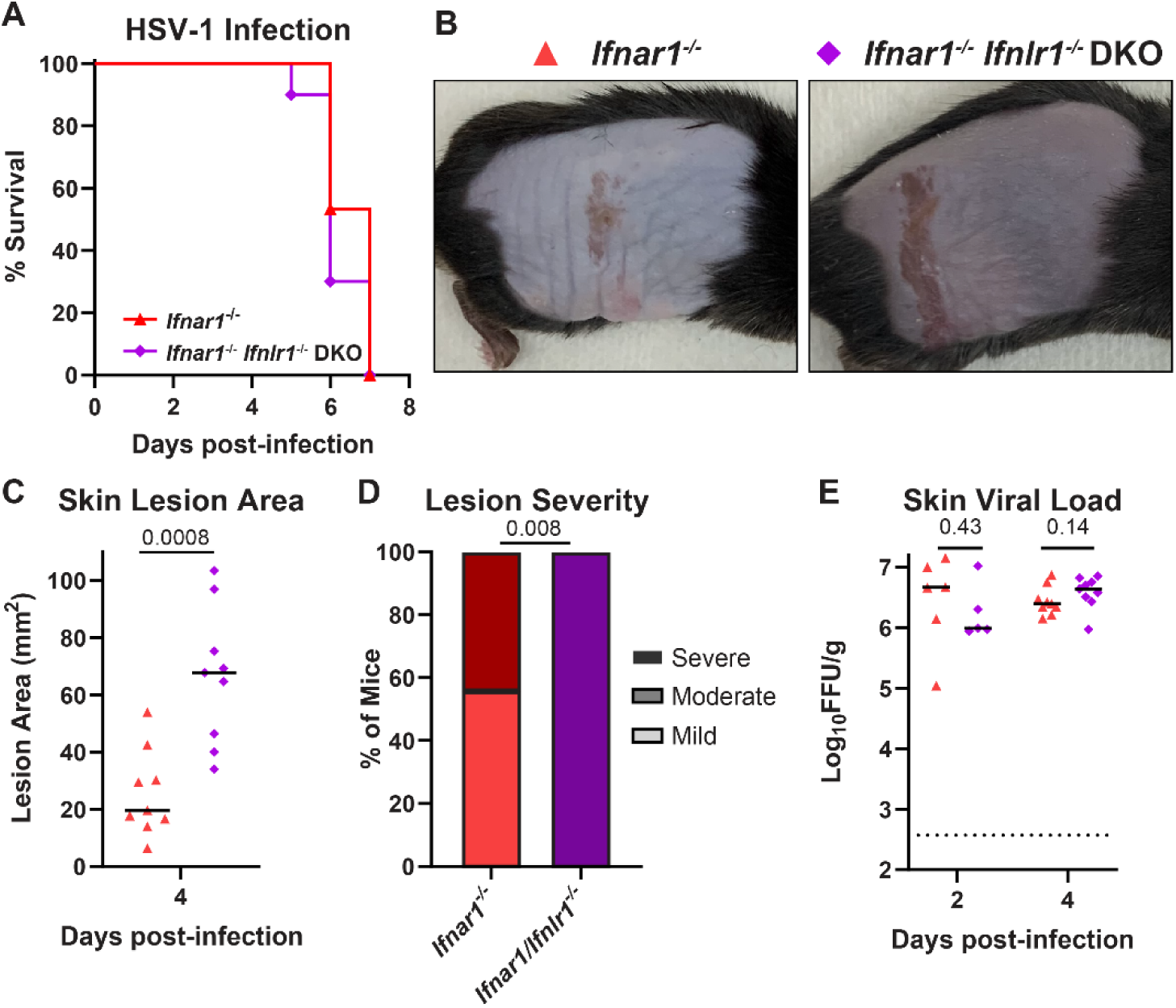
The protective effect of IFN-λ in the skin does not require IFN-α/β signaling. 8-12 week-old *Ifnar1^-/-^* and *Ifnar1^-/-^ Ifnlr1^-/-^* DKO male and female mice were infected with 10^6^ FFU of HSV-1. **A.** Lethality was monitored daily (n=15 *Ifnar1^-/-^* and 10 DKO mice). **B**. Dermatome skin lesions were photographed 4 dpi. **C.** Skin lesion areas were measured from photographs 4 dpi using ImageJ. **D.** Skin lesion severity was categorized based on 4 dpi lesion area. **E.** Skin viral loads were measured 2 and 4 dpi by focus-forming assay. Survival differences were compared by Mantel-Cox Log-Rank test. Differences in lesion area and viral load were compared by Mann-Whitney U test and differences in categorical skin disease were compared by Cochran-Armitage test. *P* values are reported with *P* < 0.05 considered to be statistically significant.

### IFN-λ cytokines protect against severe HSV-1 skin disease

We next asked which cell types produced the IFNs responsible for controlling HSV-1 skin disease. To determine whether mice lacking IFN-λ cytokines recapitulate the phenotype of mice lacking the IFN-λ receptor, we infected mice lacking IFN-λ2 and IFN-λ3 (*Ifnl2/3*^-/-^), the only IFN-λ cytokines produced in mice (Peterson et al., 2019), and measured skin lesion area, disease severity, and skin viral loads (Fig. 3A-D). At 6 dpi, *Ifnl2/3*^-/-^ mice developed significantly larger skin lesions compared to wild-type mice (median area 20.2 mm^2^ vs 7.1 mm^2^, *P* < 0.01; Fig. 3B), and a significantly higher proportion of *Ifnl2/3*^-/-^ mice developed severe skin lesions compared to wild-type mice (45% vs 15%, *P* < 0.01, Fig. 3C) with no significant difference in skin viral loads (Fig. 3D). Altogether, the phenotype of mice lacking IFN-λ cytokines was consistent with the phenotype of mice lacking the IFN-λ receptor, providing an independent line of evidence that IFN-λ signaling restricts severe HSV-1 skin disease.

**Figure 3.**
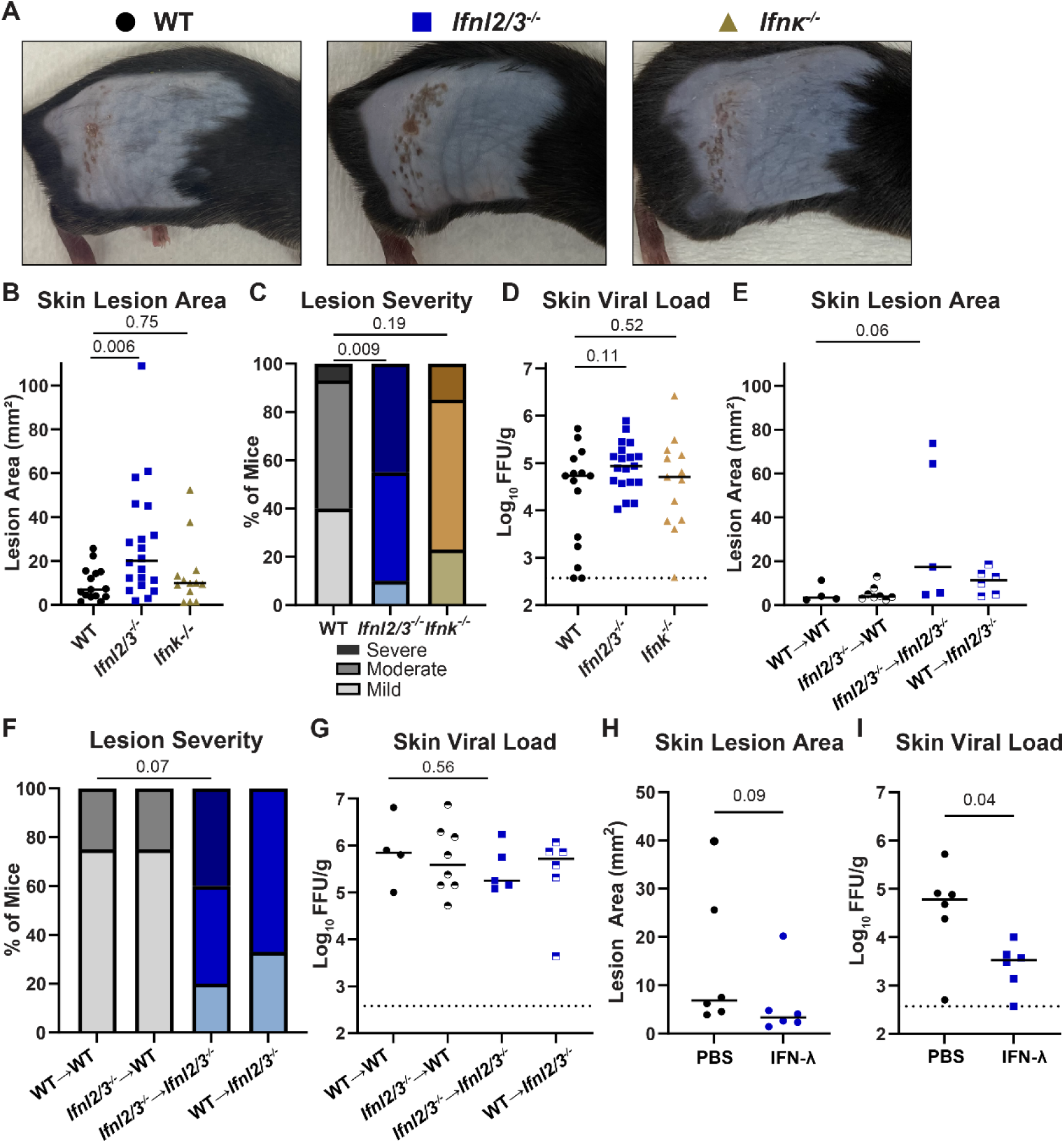
IFN-λ cytokines protect against severe HSV-1 skin disease. **A-D.** 8-12 week-old WT, *Ifnl2/3^-/-^,* and *Ifnk^-/-^* male and female mice were infected with 10^6^ FFU of HSV-1. **A**. Dermatome skin lesions were photographed 6 dpi. **B.** Skin lesion areas were measured from photographs 6 dpi using ImageJ. **C.** Skin lesion severity was categorized based on 6 dpi lesion area. **D.** Skin viral loads were measured 6 dpi by FFA. **E-G.** 8 week-old WT and *Ifnl2/3^-/-^* male and female mice were lethally irradiated and transfused with 10^7^ bone marrow cells from WT or *Ifnl2/3^-/-^* donors. 10 weeks later, mice were infected with 10^6^ FFU of HSV-1. **E.** Skin lesion areas were measured from photographs 6 dpi using ImageJ. **F.** Skin lesion severity was categorized based upon 6 dpi lesion area. **G.** Skin viral loads were measured 6 dpi by FFA. **H-I.** 8-12 week-old WT male and female mice were depilated and then topically treated with 5 µg of recombinant murine IFN-λ3 or PBS. 24 hours later, mice were infected with 10^6^ FFU of HSV-1 at the treated site. **H.** Skin lesion areas were measured from photographs 6 dpi using ImageJ. **I.** Skin viral loads were measured 6 dpi by FFA. Differences in lesion area and viral load were compared by Mann-Whitney U test and differences in categorical skin disease were compared by Cochran-Armitage test. *P* values are reported with *P* < 0.05 considered to be statistically significant.

Although some type I IFNs, such as IFN-α and IFN-β, can be produced by many cell types, IFN-κ is a type I IFN that is thought to be produced predominantly by keratinocytes (Gharaee-Kermani et al., 2022; LaFleur et al., 2001). To determine whether a keratinocyte-specific type I IFN might play a role in role in controlling skin disease caused by HSV-1, we infected *Ifnk*^-/-^ mice. We found no significant differences in lesion area, disease severity, or skin viral loads between *Ifnk*^-/-^ mice and wild-type mice (Fig. 3A-D), suggesting that, despite its keratinocyte-specific nature, IFN-κ does not play a key role controlling HSV-1 infection in the skin.

To determine the source of protective IFN-λ during HSV-1 skin infection, we used bone marrow chimeras between wild-type and *Ifnl2/3^-/-^* mice, generating mice that lacked the ability to produce IFN-λ2/3 in either their hematopoietic or stromal compartments. Due to a lack of antibodies to detect IFN-λ, we used PCR genotyping to confirm effective bone marrow engraftment in recipients. We then infected IFN-λ chimeric mice with HSV-1 and evaluated skin disease 6 dpi (Fig 3E). *Ifnl2/3*^-/-^ mice receiving *Ifnl2/3*^-/-^ bone marrow showed a trend toward larger lesions and more severe disease than wild-type mice receiving wild-type bone marrow (median area 17.3 mm^2^ vs 3.5 mm^2^, *P* = 0.06; Fig. 3E), consistent with our observations in non-irradiated mice. However, we found that mice retaining IFN-λ production in either their hematopoietic compartment or their stromal compartment were protected from severe HSV-1 skin disease (Fig. 3F), suggesting that multiple cell types can produce protective IFN-λ during HSV-1 skin infection. As in our previous experiments, the effects of IFN-λ on skin lesion severity were independent of an effect on viral load (Fig. 3G).

Exogenous IFN-λ treatment has shown therapeutic potential against various viral infections and autoimmune diseases (Blazek et al., 2015; Dinnon et al., 2020; Feld et al., 2021; Flisiak et al., 2016; Muir et al., 2014), therefore, we sought to determine whether topical administration of IFN-λ could limit HSV-1 infection and skin lesions. We depilated mice two days prior to infection, pretreated topically with 5 µg of IFN-λ and infected with HSV-1 on the treated site one day later. We found a modest decrease in lesion areas in IFN-λ pretreated mice at 6 dpi compared to PBS-pretreated mice (median area 3.4 mm^2^ vs 6.9 mm^2^, *P* = 0.09; Fig. 3G). Additionally, we found that viral loads at 6 dpi were significantly reduced in IFN-λ pretreated mice compared to PBS-pretreated mice (median viral load 3.53 Log_10_ FFU/g vs 4.78 Log_10_ FFU/g, *P* < 0.05; Fig. 3H). Altogether, these data indicate that exogenous prophylactic administration of IFN-λ can limit severe HSV-1 skin disease, suggesting potential opportunities for therapeutic interventions.

### IFN-λ signaling in both keratinocytes and leukocytes is necessary to restrict severe HSV-1 skin disease

The skin is comprised of diverse cell types including keratinocytes and resident and infiltrating leukocytes. To determine which IFN-λ responsive cell types contribute to controlling HSV-1 skin infection, we used conditional knockout mice lacking IFN-λ signaling in keratinocytes (*K14*-Cre-*Ifnlr1*^-/-^) or leukocytes (*Vav*-Cre-*Ifnlr1*^-/-^). These mice were bred as Cre hemizygotes, generating littermate Cre-controls (wild-type) and were genotyped retrospectively, allowing experiments to be performed blinded to the *Ifnlr1* status of each mouse. *Vav*-Cre-*Ifnlr1*^-/-^ mice have been previously reported (Baldridge et al., 2017; Casazza et al., 2022); however, to our knowledge this is the first derivation of *K14*-Cre-*Ifnlr1*^-/-^ mice. To validate conditional depletion of the *Ifnlr1* in these mice, we used PCR genotyping in specific tissues in wild-type (Cre-) and *K14*-Cre-*Ifnlr1*^-/-^ (Cre+) mice and found Cre-mediated rearrangement of the *Ifnlr1^f/f^* allele only in epithelial tissues (Fig. S2). Conditional knockout mice were compared to *Ifnlr1*^-/-^ mice generated by crossing *Ifnlr1*^f/f^ mice to mice expressing Cre recombinase under a CMV promoter and subsequently bred as knockout x knockout.

We infected wild-type, *Ifnlr1^-/-^*, *K14*-Cre-*Ifnlr1^-/-^*, and *Vav*-Cre-*Ifnlr1^-/-^* mice with HSV-1 and evaluated skin lesion area, disease severity, and viral loads 6 dpi (Fig. 4A-D). We found that *Ifnlr1*^-/-^ mice developed significantly larger lesions than wild-type mice (median area 13.1 mm^2^ vs 6.7 mm^2^, *P* < 0.01) (Fig. 4B) and a greater proportion of these mice developed severe skin disease (34% vs 14%, *P* < 0.01) (Fig. 4C), further supporting a role for IFN-λ in controlling severe HSV-1 skin disease in an independent line of *Ifnlr1*^-/-^ mice compared to those used in our earlier experiments (Fig. 1). We found that mice lacking IFN-λ signaling exclusively in keratinocytes or in leukocytes developed skin lesions similar in size to *Ifnlr1^-/-^* mice (median area 15.7 mm^2^, 14.6 mm^2^, and 13.1 mm^2^, *P* > 0.05) (Fig. 4C) and a similar proportion developed severe lesions (40%, 44%, 34%, *P* > 0.05) (Fig. 4D). We found no significant differences in skin viral loads comparing *Ifnlr1*^-/-^ mice to wild-type or comparing conditional knockouts to *Ifnlr1*^-/-^ (Fig. 4D). Altogether, these data indicate that IFN-λ signaling in both keratinocytes and leukocytes is necessary to control severe HSV-1 skin disease.

**Figure 4.**
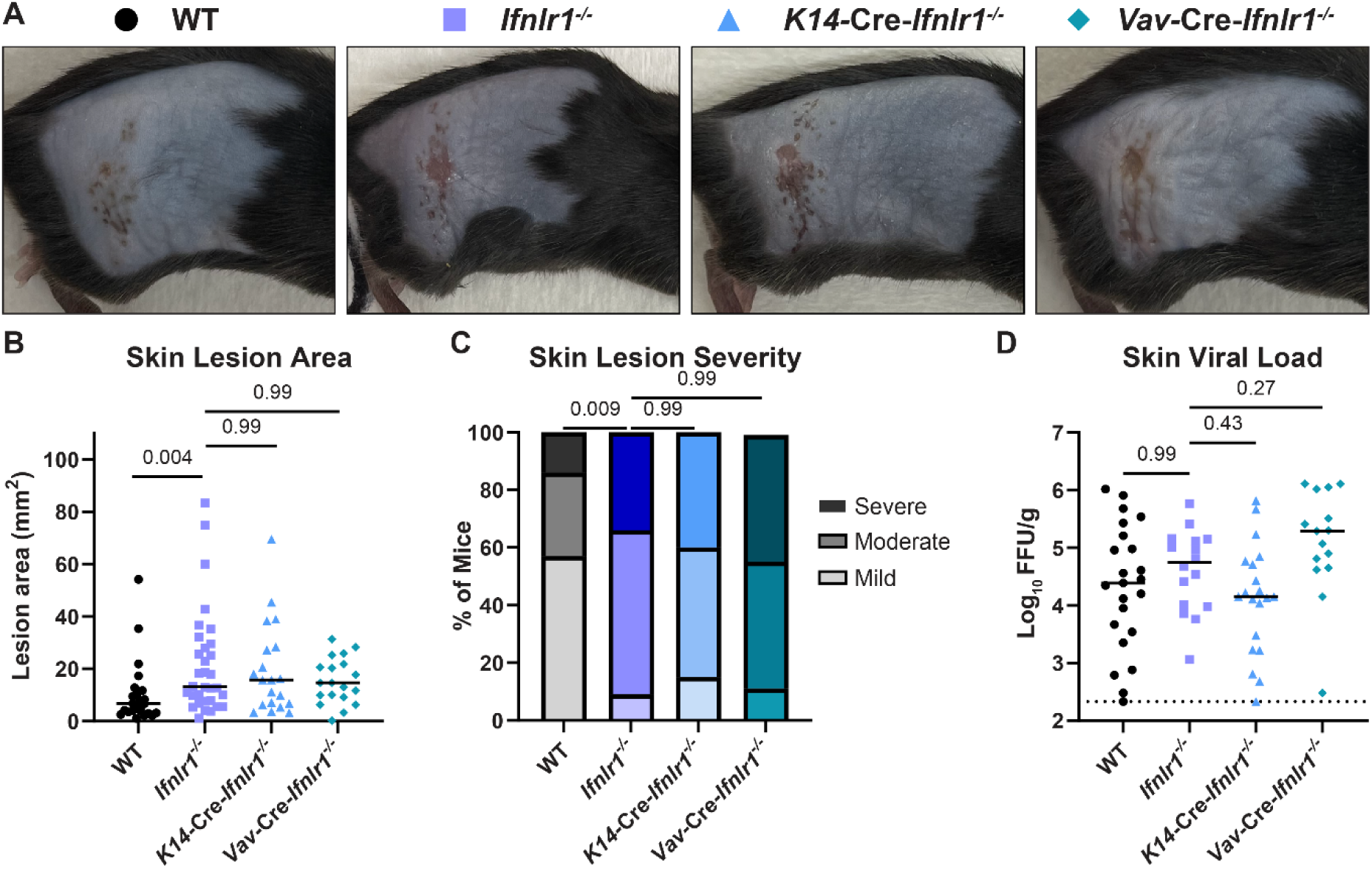
IFN-λ signaling in both keratinocytes and leukocytes is necessary to restrict severe HSV-1 skin disease. 8-12 week-old male and female WT, *Ifnlr1^-/-^, K14-*Cre*-Ifnlr1^-/-^*, and *Vav-*Cre*-Ifnlr1^-/-^* mice were infected with 10^6^ FFU of HSV-1. **A**. Dermatome skin lesions photographed were 6 dpi. **B.** Skin lesion areas were measured from photographs 6 dpi using ImageJ. **C.** Skin lesion severity was categorized based on 6 dpi lesion area. **D.** Skin viral loads were measured 6 dpi by FFA. Differences in lesion area and viral load were compared by Mann-Whitney U test and differences in categorical skin disease were compared by Cochran-Armitage test. *P* values are reported with *P* < 0.05 considered to be statistically significant.

### IFN-λ signaling in keratinocytes differentially regulates inflammatory genes, including CXCL9

Because we found that IFN-λ signaling in both keratinocytes and leukocytes was necessary to control HSV-1 skin lesion severity, we next asked whether this was due to crosstalk between keratinocytes and leukocytes. To understand how IFN-λ could be mediating signaling between epithelial cells and immune cells, we first generated IFN receptor knockout A549 cells, a human lung epithelial cell line. We treated *IFNLR1* KO, *IFNAR1* KO, or nontargeting control (RenLuc) A549 cells with recombinant IFN-β or IFN-λ and measured induction of the ISG *IFIT1* by qRT-PCR. As expected, *IFNLR1* KO cells responded to IFN-β but not IFN-λ and *IFNAR1* KO cells responded to IFN-λ but not IFN-β (Fig. S4A). We then treated control and IFN receptor KO cells with 50 ng/mL IFN-λ, 5 ng/mL IFN-β, or media alone for 8 hours and performed bulk RNAseq (Fig. S4B-F). We found that the transcriptional response after IFN-λ treatment in A549 cells was modest compared to IFN-β (Supplementary Table 1), consistent with prior studies (Coldbeck-Shackley et al., 2023; Zhou et al., 2007). However, prior studies that used primary cells rather than cell lines found more robust transcriptional responses induced by IFN-λ (Caine et al., 2019; Galani et al., 2017). We therefore investigated IFN-λ induced transcriptional responses in primary keratinocytes.

To define the IFN-λ specific response in keratinocytes and understand how IFN-λ could be signaling through keratinocytes to mediate their crosstalk with leukocytes, we generated primary keratinocytes from wild-type, *Ifnar1*^-/-^, *Ifnlr1*^-/-^, and *Ifnar1*^-/-^ *Ifnlr1*^-/-^ DKO mice, and treated them with 5 ng IFN-λ, 5 ng IFN-β, or media for 24 hours. We first measured induction of *Ifit1* by qRT-PCR. As expected, IFN-β did not induce a response in *Ifnar1*^-/-^ cells, IFN-λ did not induce a response in *Ifnlr1*^-/-^ cells, and neither IFN induced a response in *Ifnar1*^-/-^ *Ifnlr1*^-/-^ cells (Fig. 5A). We then performed bulk RNAseq with the goal of identifying IFN-λ-specific antiviral inflammatory mediators. By principal component analysis, triplicate samples clustered together, but different genotypes were distinct even without IFN treatment, suggesting differences in tonic IFN signaling in keratinocytes (Fig 5B). While the IFN-β response was largely unchanged in *Ifnlr1*^-/-^ cells compared to wild-type, the IFN-λ response was diminished in *Ifnar1*^-/-^ cells by both qRT-PCR (Fig. 5A) and principal component analysis (Fig. 5B), suggesting cross-talk between IFN-λ and IFN-β signaling in keratinocytes as well as in A549 cells (Fig. S4B). Although the IFN-λ response generally is reported to be less potent than the IFN-α/β response (Fig. S4) (Lazear et al., 2019), we found that many ISGs (including *Ifit1* and *Rsad2*) were induced to a similar magnitude by IFN-λ and IFN-β (Fig. 5A, C). We found a subset of ISGs in primary keratinocytes to be differentially induced by IFN-λ but not IFN-β (Fig. 5D). However, upon inspecting the IFN-λ-specific gene list we found that while 11 or 10 genes did meet our differential threshold (2-fold change, FDR *P* < 0.05), these genes all were induced to a similar extent by IFN-β (although not quite reaching the differential threshold) (Fig. 5E). Moreover, none of the genes identified as IFN-λ-specific in wild-type or knockout keratinocytes overlapped (Supplementary Table 2), arguing against a set of ISGs that are induced in an IFN-λ specific manner in this system.

**Figure 5.**
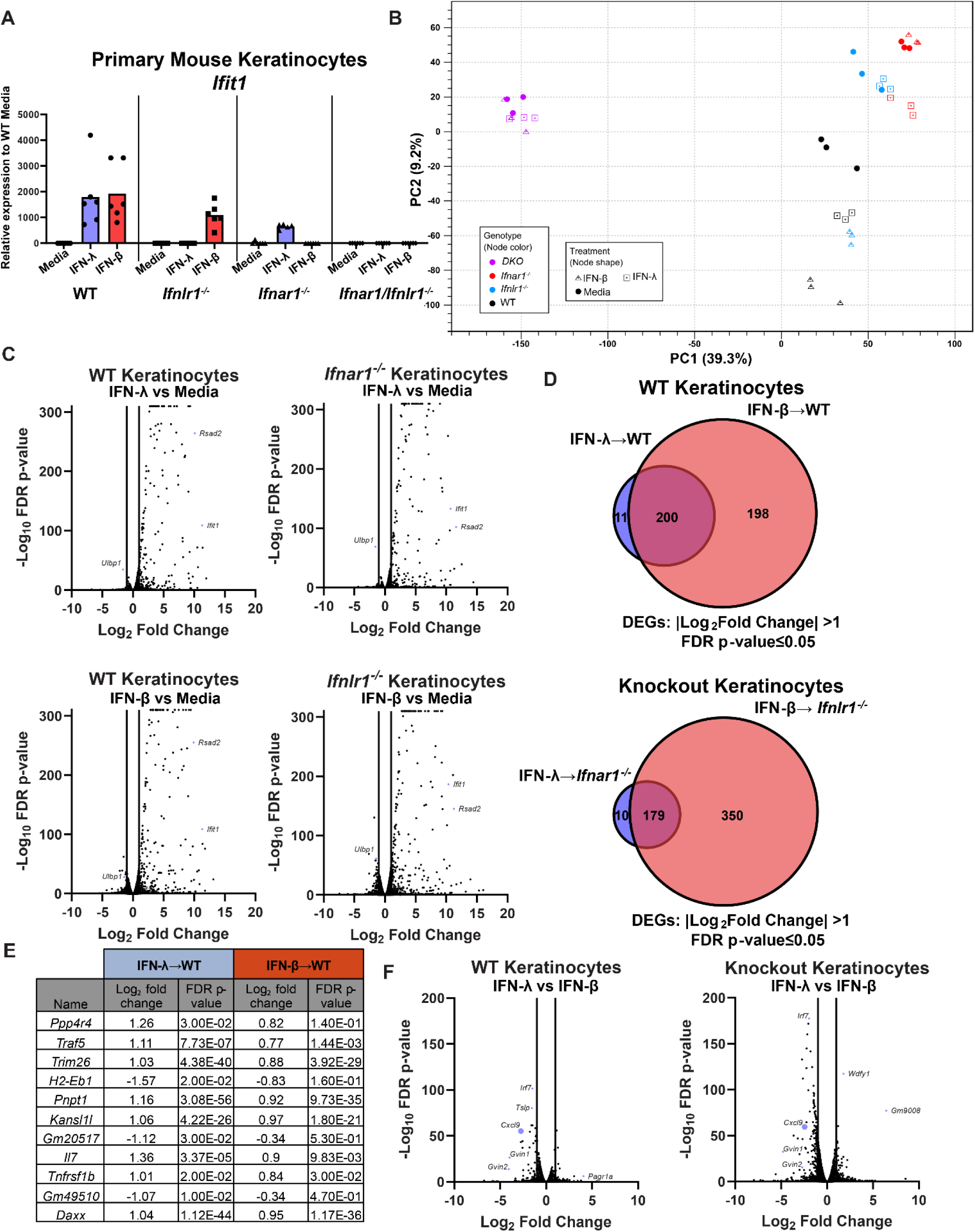
IFN-λ signaling in keratinocytes differentially regulates CXCL9 induction. Primary keratinocytes were prepared from WT, *Ifnlr1*^-/-^, *Ifnar1*^-/-^, and *Ifnar1*^-/-^ *Ifnlr1*^-/-^ DKO mice and treated with IFN-λ3 (5 ng/mL), IFN-β (5 ng/mL), or media alone for 24 hours. **A.** RNA was extracted and IFN-stimulated gene expression was measured by qRT-PCR. *Ifit1* expression is shown relative to *ActB* (housekeeping gene) and normalized to expression in media-treated WT cells. Results represent 6 samples from 2 independent experiments. **B-F.** RNA from 3 samples per genotype per treatment was analyzed by RNAseq (NovaSeq6000S4 XP Paired End 2×100). **B.** Principal component analysis for all analyzed samples. **C.** Volcano plots showing differentially expressed genes after IFN-λ or IFN-β treatment, compared to media-only treated keratinocytes. Differentially expressed genes were defined as having a |Log_2_Fold Change| >1 and an FDR p-value ≤ 0.05. **D.** Venn diagram showing DEGs induced by IFN-λ and IFN-β in WT or knockout keratinocytes. **E.** Table of IFN-λ-specific DEGs, their Log_2_Fold Change, and FDR p-value for IFN-λ and IFN-β-treated WT keratinocytes. **F.** Volcano plot showing DEGs after IFN-λ treatment compared to IFN-β treatment in WT and receptor knockout keratinocytes.

We next analyzed the IFN-λ response relative to the IFN-β response in WT and knockout keratinocytes to identify genes that are differentially regulated by IFN-λ compared to IFN-β (Fig. 5F). We found that the chemokine CXCL9 was one of the top differentially regulated genes between IFN-λ and IFN-β treatment in both wild-type and the corresponding receptor knockout keratinocytes. Notably, *Cxcl9* was significantly less induced after IFN-λ treatment compared to IFN-β in wild-type keratinocytes (9.7-fold vs 65.0-fold) and knockout keratinocytes (23.0-fold vs 58.5-fold). Because *Cxcl9* is known to recruit a variety of immune cells such as T cells, NK cells, and neutrophils (Chami et al., 2014, 2017; Karin, 2020), we next sought to characterize IFN-λ-dependent leukocyte changes in HSV-1 infected skin to identify possible players in IFN-λ-mediated protection by keratinocytes and leukocytes.

### Skin lesions in Ifnlr1^-/-^ mice exhibit extensive neutrophil infiltration and severe pathology

To define IFN-λ dependent changes in leukocyte populations in response to HSV-1 infection, we harvested skin lesions and adjacent healthy skin at 6 dpi from *Ifnlr1*^-/-^ and wild-type mice and analyzed leukocyte populations by flow cytometry. We found no significant difference between *Ifnlr1*^-/-^ and wild-type leukocyte population frequencies in the healthy skin at 6 dpi (Fig. S3A). Further, we found no differences in leukocyte frequency, including dendritic cells, macrophages, CD4 and CD8 T cells, B cells, or NK cells, between *Ifnlr1*^-/-^ and wild-type skin lesions at 6 dpi (Fig. S3A). However, we found that skin lesions from *Ifnlr1*^-/-^ mice exhibited a significantly higher frequency of neutrophils (CD45+CD11b+Ly6G+, Fig. 6A-B) compared to wild-type mice at 6 dpi (mean neutrophil frequency 21.1% vs 6.7%; Fig. 6B). Although we found a significant increase in γδT cells between *Ifnlr1*^-/-^ and wild-type mice (Fig. S3A), we did not further investigate these cells as they are not known to be IFN-λ responsive and are less abundant in the skin compared to neutrophils. This increase in neutrophil frequency was specific to the skin lesion, because in adjacent healthy skin neutrophil abundance was very low and there was no significant difference between *Ifnlr1*^-/-^ and wild-type mice (Fig. 6B). The increase in neutrophil frequency in *Ifnlr1*^-/-^ mice corresponded to the kinetics of skin lesion formation, as overall neutrophil frequency was low at 4 dpi, with no significant difference between wild-type and *Ifnlr1^-/-^* mice and the increased neutrophil frequency in *Ifnlr1*^-/-^ compared to wild-type mice was first evident during peak disease at 6 dpi and maintained through 8 dpi (Fig. 6B).

**Figure 6.**
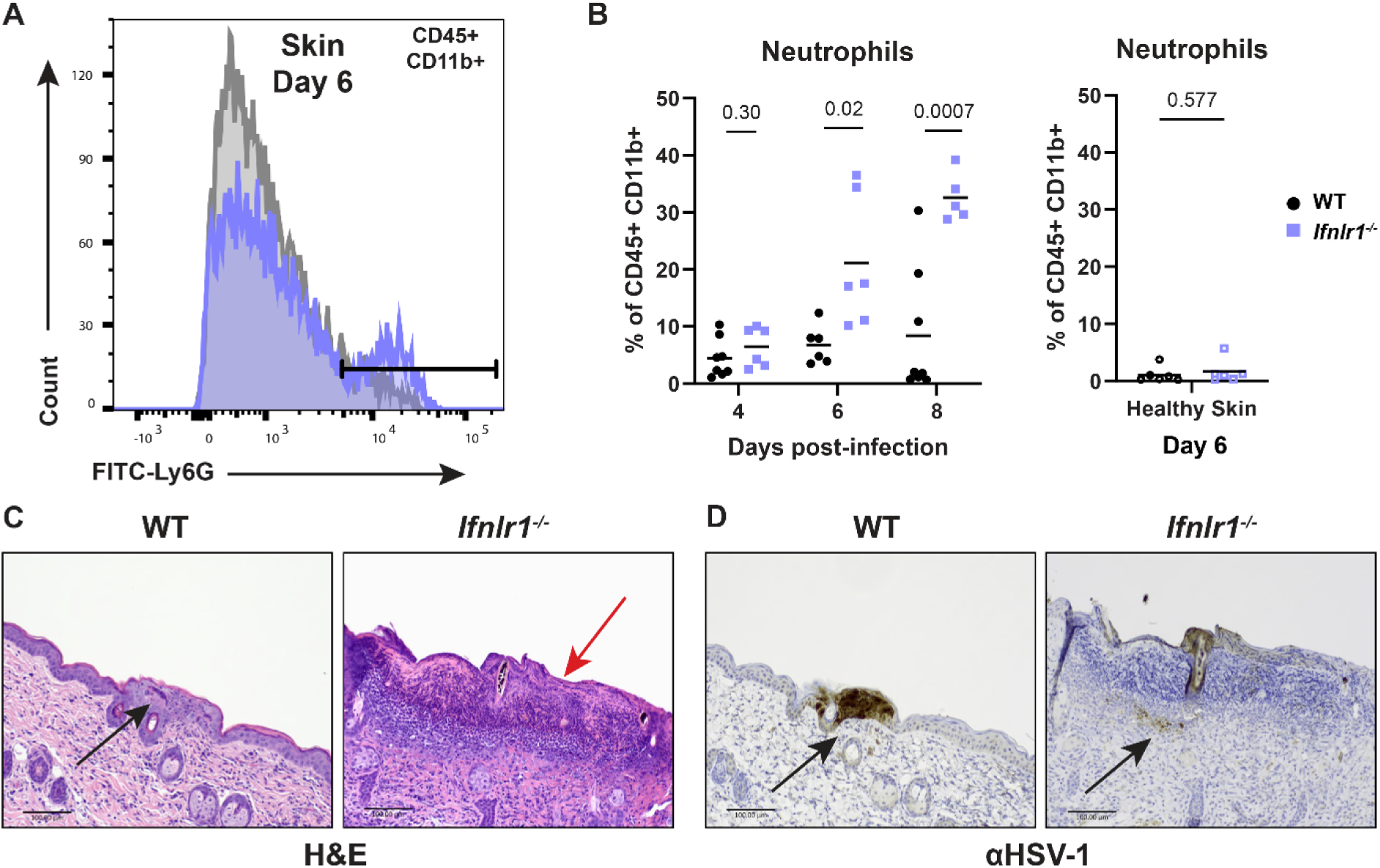
IFN-λ signaling limits neutrophil abundance and severe skin pathology following HSV-1 infection. 8-12 week-old male and female WT and *Ifnlr1^-/-^* mice were infected with 10^6^ FFU of HSV-1. **A-B.** Dermatome lesions and adjacent healthy skin were collected 4-8 dpi and analyzed by flow cytometry. **A.** Representative histogram of WT and *Ifnlr1^-/-^* lesions showing Ly6G+ neutrophil gating strategy (CD45+, CD11b+, Ly6G+) at 6 dpi. **B.** Frequency of neutrophils (Ly6G+ out of CD45+ CD11b+ live cells) for WT and *Ifnlr1^-/-^* dermatome lesions and adjacent healthy skin. Significant differences in neutrophil frequency were determined using an unpaired t test. *P* values are reported with *P* < 0.05 considered to be statistically significant. **C-D.** Inoculated flank skin was collected at 6 dpi and serial sections of the same skin lesion were processed for histology. **C.** H&E staining; black arrow denotes a neutrophilic pustule, red arrow denotes diffuse neutrophilic infiltrate. **D.** Anti-HSV-1 immunohistochemistry; viral antigen staining is denoted by black arrows. Scale bars are 100 µm.

To better define the spatial organization of IFN-λ mediated control of neutrophil infiltration, we harvested skin lesions and healthy skin 6 dpi from wild-type and *Ifnlr1*^-/-^ mice and evaluated skin pathology by H&E staining (Fig. 6C) and immunohistochemistry for HSV-1 antigen (Fig. 6D). We found that *Ifnlr1*^-/-^ mice displayed more significant skin damage and overall loss or necrosis of the epidermis in their skin lesions compared to wild-type mice (Fig. 6C, red arrow). In contrast, wild-type mice had a greater number of intact pustules (Fig. 6C, black arrow) compared to *Ifnlr1*^-/-^ mice, which had more diffuse spread of neutrophils throughout the dermis and epidermis. Further, wild-type mice also had an increased staining intensity for HSV-1 antigen compared to *Ifnlr1*^-/-^ mice, in which most staining was evident within pustules (Fig. 6D). HSV-1 antigen was more present in the dermis for *Ifnlr1*^-/-^ mice (black arrows). However, the epidermis was often missing or necrotic in many areas of intense neutrophilic inflammation in *Ifnlr1*^-/-^ mice, possibly precluding antigen detection in the epidermis. Altogether, these data suggest that a lack of IFN-λ signaling results in greater neutrophil infiltration and more severe HSV-1 skin pathology.

### IFN-λ signaling suppresses neutrophil-mediated pathology to limit HSV-1 skin disease

To further define the role of neutrophils in driving HSV-1 skin lesion pathology in the absence of IFN-λ signaling, we depleted neutrophils from wild-type and *Ifnlr1^-/-^* mice 0, 2, and 4 dpi using a αLy6G-depleting antibody (or an isotype-control antibody or PBS) and evaluated dermatome skin lesions 6 dpi (Fig. 7A). To confirm depletion of circulating neutrophils, we collected splenocytes 6 dpi (Fig. 7B) and detected neutrophils (Ly6G+) by flow cytometry. We found that αLy6G antibody efficiently depleted neutrophils compared to isotype-control treated mice (9.8% vs 0.8% Ly6G+ of CD11b+) (Fig. 7B). We found that neutrophil depletion had no effect on skin lesion area in wild-type mice, consistent with prior studies showing that neutrophils do not mediate HSV-1 skin lesion pathology in wild-type mice (Wojtasiak et al., 2010). However, we found that while *Ifnlr1^-/-^* mice developed significantly larger lesions than wild-type mice and a greater proportion developed severe lesions in mice with intact neutrophils (PBS or isotype-control), IFN-λ-dependent protection was lost in neutrophil-depleted mice (Fig. 7C-D). Altogether these results suggest that IFN-λ protects against severe HSV-1 skin disease by suppressing neutrophil-driven immunopathology.

**Figure 7.**
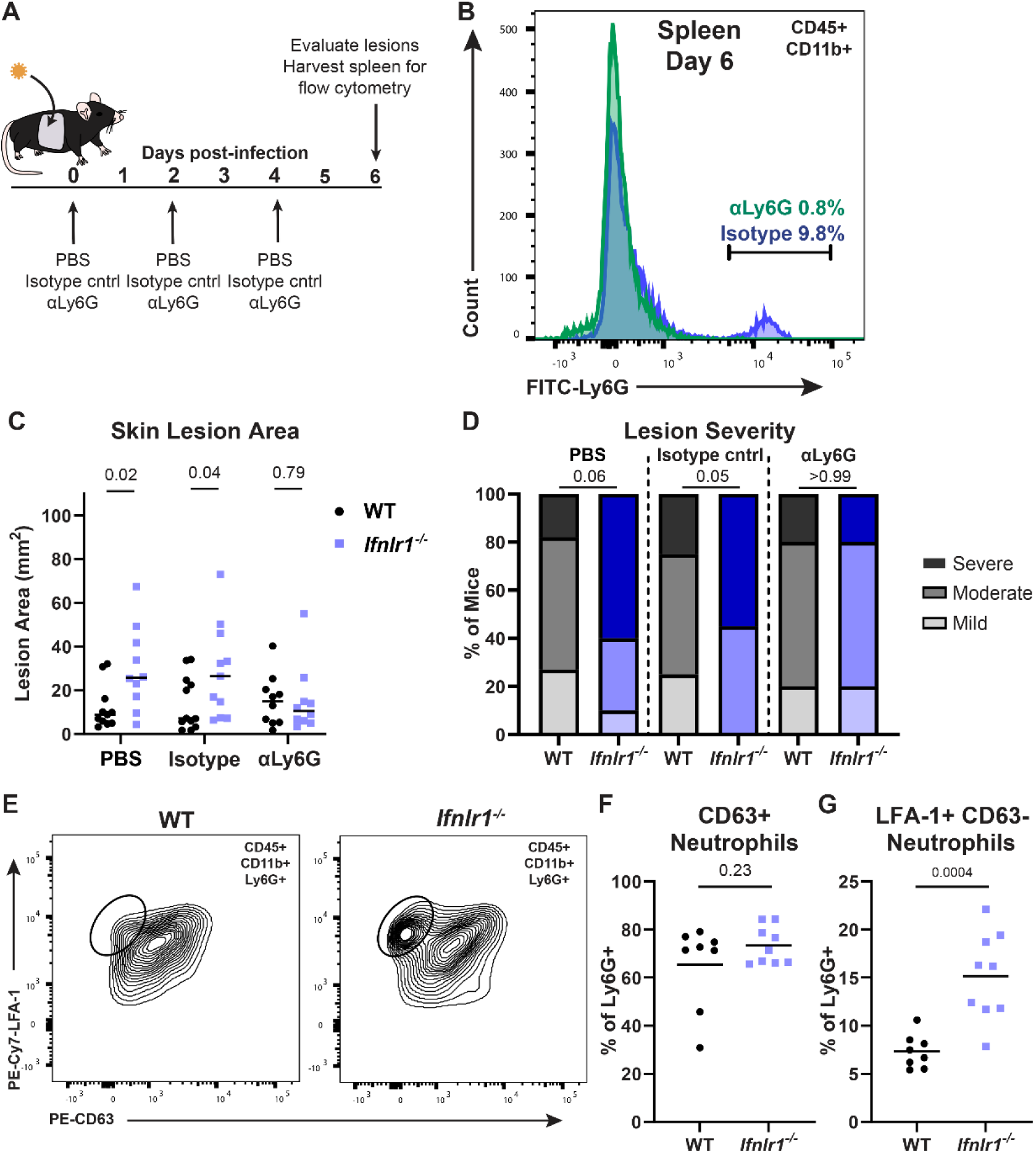
IFN-λ signaling suppresses neutrophil-mediated pathology to limit HSV-1 skin disease. **A.** Experimental design for neutrophil depletions **B-D**. WT and *Ifnlr1^-/-^* mice were infected with 10^6^ FFU of HSV-1. 0, 2, and 4 dpi mice were injected intraperitoneally with PBS, 250 µg of αIsotype control (IgG2a), or 250 µg of αLy6G (Clone 1A8). **B.** Spleens were harvested 6 dpi and analyzed by flow cytometry to confirm neutrophil depletion. **C.** Skin lesions were photographed 6 dpi and lesion areas measured ImageJ. **D.** Skin lesion severity was categorized based on 6 dpi lesion area. **E-G.** WT and *Ifnlr1^-/-^* mice were infected with 10^6^ FFU of HSV-1 and 6 dpi skin lesions were photographed and analyzed via flow cytometry. **E.** Representative plots and gating strategy for neutrophil phenotyping markers LFA-1 and CD63 (% of CD45+ CD11b+ Ly6G+). **F.** Frequency of CD63+ neutrophils. **G.** Frequency of LFA-1+ CD63-neutrophils. Differences in lesion area and viral load were compared by Mann-Whitney U test and differences in categorical skin disease were compared by Cochran-Armitage test. Neutrophil frequencies were compared by unpaired t test. *P* values are reported with *P* < 0.05 considered to be statistically significant.

To determine the mechanism by which IFN-λ suppresses neutrophil-mediated pathology during HSV-1 skin infection, we investigated whether IFN-λ signaling impacts neutrophil recruitment and/or effector function. We infected wild-type and *Ifnlr1*^-/-^ mice with HSV-1 and collected skin lesions at 6 dpi for flow cytometry. We measured neutrophil expression of lymphocyte function associated antigen 1 (LFA-1, an integrin upregulated upon neutrophil activation and associated with neutrophil recruitment from circulation into inflamed tissues) and CD63 (a lysosomal membrane protein expressed on primary neutrophil granules that is presented on the plasma membrane upon the final stage of neutrophil degranulation) (Fig. 7E-G) (Eichelberger et al., 2019; Phillipson et al., 2006; Zenaro et al., 2015). We found no significant difference in primary granule release (% LFA-1+CD63+) between wild-type and *Ifnlr1^-/-^* mice (65.4% vs 73.3%, *P* > 0.05) (Fig. 7F). However, we found that skin lesions from *Ifnlr1^-/-^* mice had a greater proportion of LFA-1+ CD63-neutrophils than wild-type mice, indicative of a population of neutrophils in the skin that are activated and accumulating but have not yet fully degranulated (15.2% vs 7.3%, *P* < 0.05; Fig. 7G). Therefore, our data support a model in which IFN-λ signaling suppresses the accumulation, but not effector functions, of neutrophils in HSV-1 skin lesions, protecting against the development of severe skin pathology.

Neutrophils are known to respond to IFN-λ in the context of other infection and inflammation models in mice (Blazek et al., 2015; Broggi et al., 2017; Espinosa et al., 2017), so we next asked whether the protective effects of IFN-λ against neutrophil-mediated pathology in in the skin required IFN-λ signaling directly in neutrophils. We infected conditional knockout mice lacking IFN-λ signaling in myeloid cells (*LysM*-Cre-*Ifnlr1*^-/-^) or neutrophils (*Mrp8*-Cre-*Ifnlr1*^-/-^) with HSV-1 and evaluated skin lesions and disease severity at 6 dpi, compared to wild-type and *Ifnlr1^-/-^* mice (Fig. 8A-C). We found that mice that lack IFN-λ signaling exclusively in all myeloid cells or specifically in neutrophils developed skin lesions similar in size to *Ifnlr1^-/-^* mice (mean area 21.5 mm^2^, 19.4 mm^2^, and 25.0 mm^2^, *P* > 0.05) (Fig. 8B) and with a similar proportion developing severe lesions (45%, 27%, 58%, *P* > 0.05) (Fig. 8C). Altogether, these data support a model where IFN-λ signals through keratinocytes and myeloid cells, including neutrophils, to suppress neutrophil accumulation in the skin during HSV-1 infection to control severe HSV-1 skin disease.

**Figure 8.**
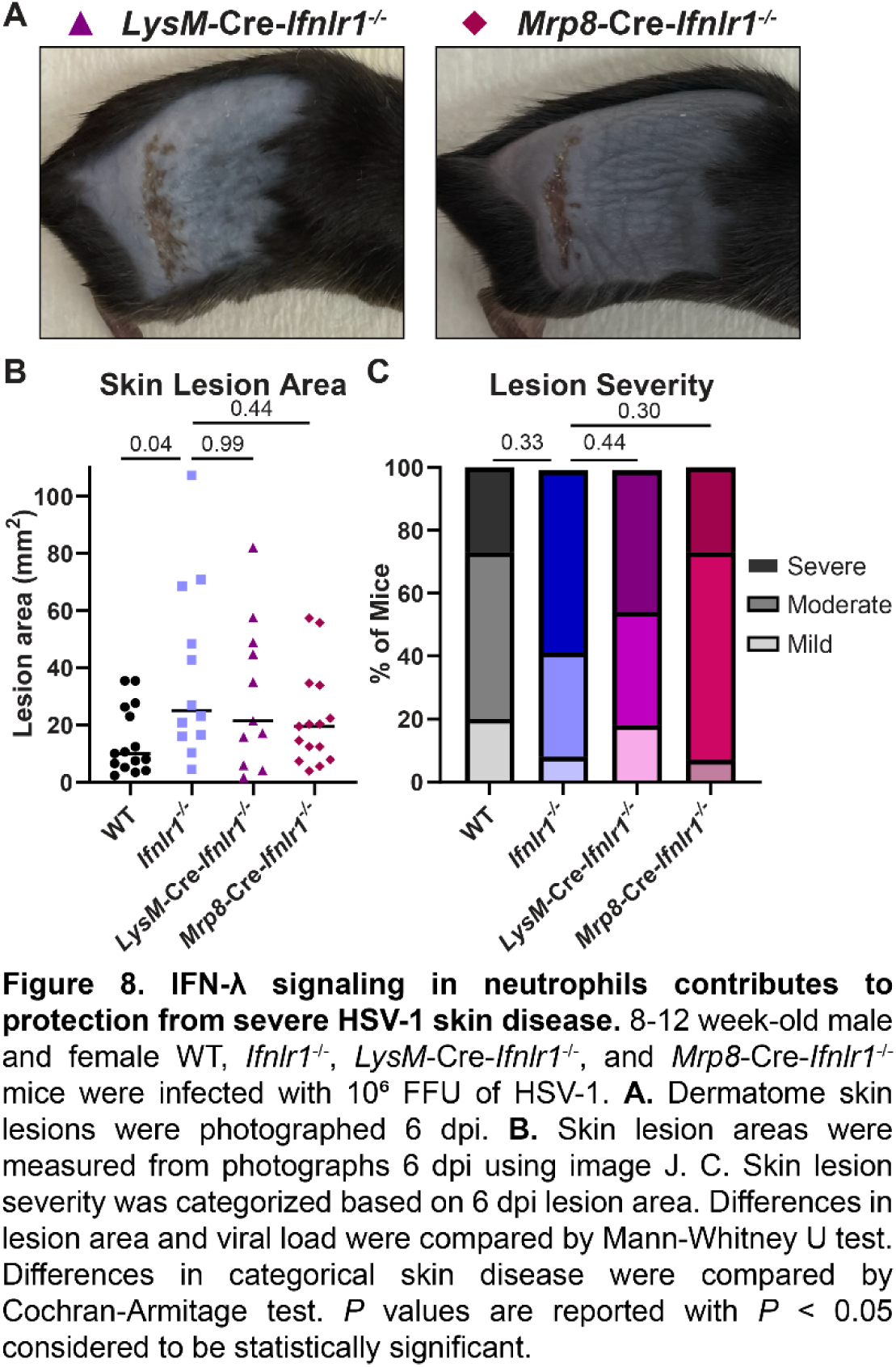
IFN-λ signaling in neutrophils contributes to protection from severe HSV-1 skin disease. 8-12 week-old male and female WT, *Ifnlr1^-/-^, LysM-*Cre*-Ifnlr1^-/-^*, and *Mrp8-*Cre*-Ifnlr1^-/-^* mice were infected with 10^6^ FFU of HSV-1. **A**. Dermatome skin lesions were photographed 6 dpi. **B.** Skin lesion areas were measured from photographs 6 dpi using image J. **C.** Skin lesion severity was categorized based on 6 dpi lesion area. Differences in lesion area and viral load were compared by Mann-Whitney U test. Differences in categorical skin disease were compared by Cochran-Armitage test. *P* values are reported with *P* < 0.05 considered to be statistically significant.

### IFN-λ signaling in keratinocytes regulates CXCL9 production to restrict neutrophil infiltration and HSV-1 skin pathology

Having established neutrophil recruitment as a key mechanism of IFN-λ-mediated protection against HSV-1 skin disease, we next sought to determine whether the chemokine CXCL9, identified as a DEG in IFN-λ-treated primary keratinocytes (Fig. 5F), mediated neutrophil recruitment in the absence of IFN-λ signaling. Although classically a lymphocyte chemoattractant, neutrophils also are reported express CXCR3, the receptor for CXCL9, in inflammatory contexts (Boff et al., 2018, 2022; Chami et al., 2014, 2017; Hartl et al., 2008). Furthermore, Forero et al. previously reported that IFN-λ failed to induce expression of *Cxcl9* and other chemokines downstream of a lack of IRF1 induction (Forero et al., 2019). Therefore, we next asked whether IFN-λ acts though keratinocytes via CXCL9 to limit neutrophil infiltration and severe HSV-1 skin disease. First, we measured the concentration of CXCL9 in dermatome lesions and adjacent healthy skin of wild-type and *Ifnlr1^-/-^* mice 6 dpi (Fig. 9A). We found that *Ifnlr1^-/-^* mice had significantly higher CXCL9 concentrations in their skin lesions compared to wild-type mice (mean 3191 pg/g vs 1634 pg/g, *P* = 0.02). The suppressive effect of IFN-λ on CXCL9 was specific to lesioned skin because in adjacent healthy skin CXCL9 concentrations were low and there was no significant difference between wild-type and *Ifnlr1^-/-^* mice.

**Figure 9.**
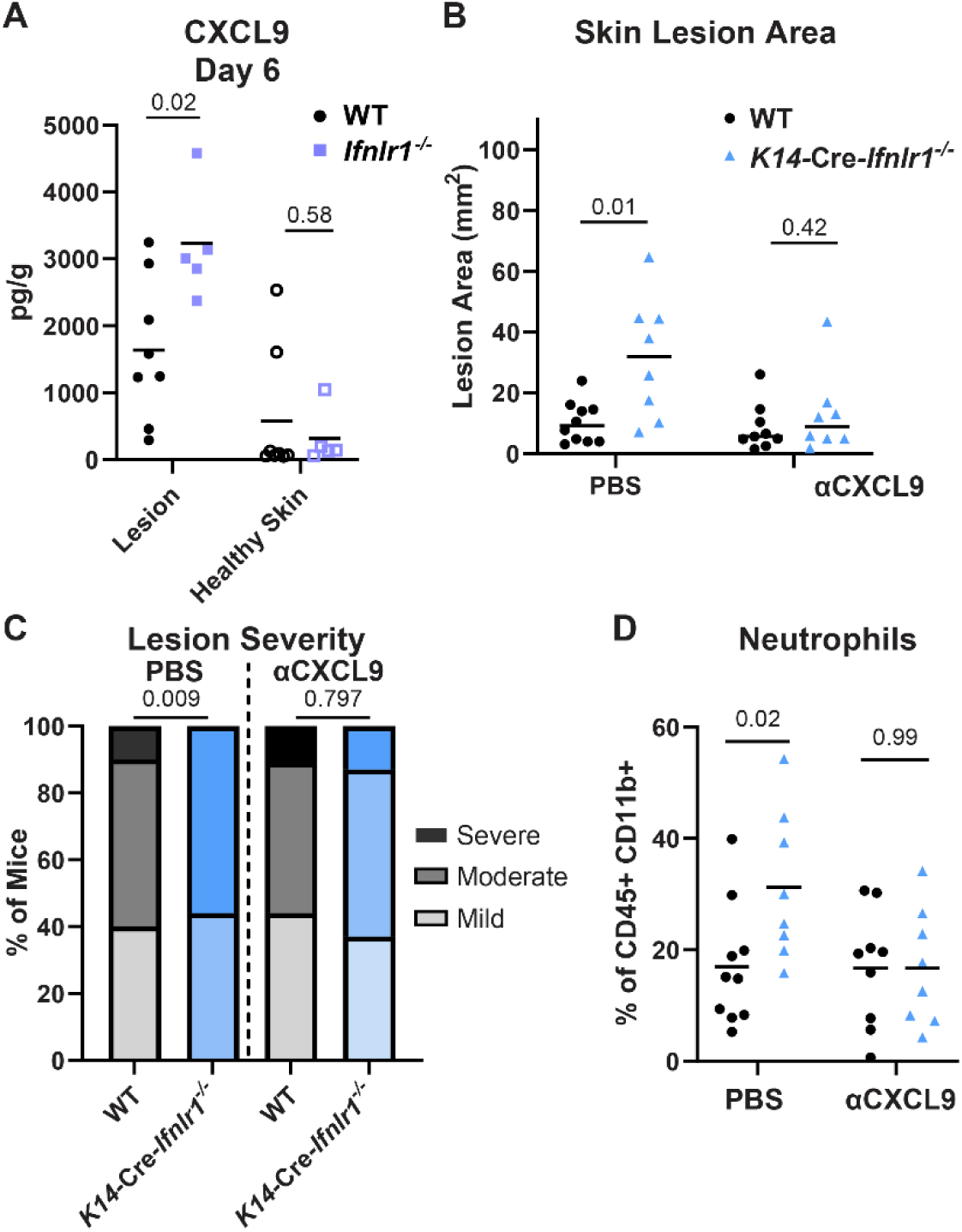
IFN-λ signaling in keratinocytes restricts CXCL9 production to limit neutrophil recruitment and HSV-1 skin disease. 8-12 week-old male and female mice were infected with 10^6^ FFU of HSV-1 and skin lesions were analyzed 6 dpi. **A.** CXCL9 was measured by ELISA in homogenates of lesional skin or adjacent healthy skin from WT and *Ifnlr1^-/-^* mice. **B-D.** WT and *K14-*Cre*-Ifnlr1^-/-^* mice were infected with HSV-1 and injected intraperitoneally with anti-CXCL9 (300µg) or PBS 0, 2, and 4 dpi. Skin lesion areas were measured from photographs 6 dpi (**B**) and lesion severity categorized (**C**). **D.** Neutrophil frequency in skin lesions was measured by flow cytometry 6 dpi. Differences in lesion area and viral load were compared by Mann-Whitney U test. Differences in categorical skin disease were compared by Cochran-Armitage test. Differences in CXCL9 concentration and neutrophil frequency were compared by unpaired t test. *P* values are reported with *P* < 0.05 considered to be statistically significant.

Next, we asked whether CXCL9 contributed to neutrophil recruitment and skin pathology, and whether this occurred downstream of IFN-λ signaling in keratinocytes. We depleted CXCL9 by injecting wild-type and *K14*-Cre-*Ifnlr1*^-/-^ mice with CXCL9 neutralizing antibody or PBS intraperitoneally 0, 2, and 4 dpi and assessed lesion area, disease severity, and skin neutrophil populations by flow cytometry (Fig. 9B-D). We did not observe any effect of αCXCL9 treatment on lesion area, disease severity, or neutrophil skin infiltration in wild-type mice. Consistent with prior experiments (Fig. 4), we found that PBS-treated *K14*-Cre-*Ifnlr1*^-/-^ mice had larger lesions and more severe disease than wild-type mice (median lesion area 13.0 mm^2^ vs 6.7 mm^2^, *P* = 0.01). We also found that *K14*-Cre-*Ifnlr1*^-/-^ mice had significantly greater neutrophil skin infiltration compared to wild-type mice (mean neutrophil frequency 31.3% vs 16.9%, P = 0.02). Strikingly, neutralizing CXCL9 ablated these differences, with *K14*-Cre-*Ifnlr1*^-/-^ mice exhibiting similar lesion areas, disease severity, and neutrophil infiltration compared to wild-type mice after αCXCL9 treatment (Fig.9B-D). Altogether, these data suggest that IFN-λ regulates CXCL9 production in keratinocytes which restricts neutrophil skin infiltration and severe HSV-1 skin disease.

Overall, we report a new protective role for IFN-λ signaling against HSV-1 skin infection. We found that IFN-λ signaling in both keratinocytes and neutrophils is necessary to mediate protection from severe HSV-1 skin disease. We uncovered a mechanism by which IFN-λ signaling in keratinocytes suppresses CXCL9 production to control neutrophil-mediated pathology by limiting neutrophil recruitment and retention in the skin. This is the first report of a protective role for IFN-λ signaling in the skin against an infection and suggests future opportunities for therapeutic uses of IFN-λ against HSV-1 and other skin pathogens.

## DISCUSSION

IFN-λ elicits antiviral immunity and maintains barrier integrity at epithelial surfaces, controlling infections locally without the inflammatory immune pathology triggered by the more potent systemic IFN-αβ response (Lazear et al., 2019). Although IFN-λ induces an antiviral transcriptional program most potently in epithelial cells, IFN-λ signaling in leukocytes, such as neutrophils and dendritic cells (DCs), is key to the ability of IFN-λ to induce a protective response without damaging inflammation (Broggi et al., 2017; Espinosa et al., 2017; Galani et al., 2017; Hemann et al., 2019). Despite the well-established role for IFN-λ in antiviral immunity at anatomic barriers (Lazear et al., 2019), including the respiratory and gastrointestinal tracts (Grau et al., 2020; Peterson et al., 2019; Ye et al., 2019), the blood-brain barrier (Douam et al., 2017; Lazear et al., 2015), and the maternal-fetal interface (Casazza et al., 2022; Jagger et al., 2017), the effects of IFN-λ in the skin have not been extensively investigated. To investigate the role of IFN-λ signaling in the skin, we used HSV-1, an important human pathogen that targets epithelial surfaces including the skin, and which has a well-established mouse model for skin infection (Simmons & Nash, 1984; van Lint et al., 2004; Wang et al., 2019). We found that mice lacking IFN-λ signaling (*Ifnlr1*^-/-^, *Ifnl2/3*^-/-^, or *Ifnar1*^-/-^ *Ifnlr1*^-/-^) developed larger HSV-1 skin lesions compared to their IFN-λ sufficient counterparts. However, lesion size was uncoupled from skin viral loads, suggesting a role for IFN-λ in suppressing inflammatory immune pathology during HSV-1 skin infection. Using conditional knockout mice, we found that IFN-λ signaling in both keratinocytes and leukocytes was necessary for protection from severe skin disease. We further showed that neutrophils are required for the enhanced skin pathology observed in *Ifnlr1*^-/-^ mice and that IFN-λ signaling in neutrophils contributes to the protective effect of IFN-λ against HSV-1 skin disease. Altogether, our results support a model where IFN-λ exerts an immunomodulatory role during HSV-1 skin infection, suppressing neutrophil-mediated pathology.

IFN-λ suppresses neutrophil recruitment and pathology in colitis and rheumatoid arthritis models (Blazek et al., 2015; Broggi et al., 2017), as well as during influenza virus infection (Galani et al., 2017). Our results add to a growing number of systems that demonstrate a role for neutrophils as a key IFN-λ responsive cell type in mice and broaden the range of mechanisms by which IFN-λ impacts neutrophil function, although there is still debate regarding the extent to which human neutrophils respond to IFN-λ (Broggi et al., 2017; Espinosa et al., 2017; Goel et al., 2020; Santer et al., 2020). Neutrophils are recruited to the skin during HSV-1 skin infection but are not important for mediating skin pathology in wild-type mice (Hor et al., 2017; Wojtasiak et al., 2010). Our studies revealed a pathogenic role for neutrophils during HSV-1 skin infection but only in the absence of IFN-λ signaling, identifying a key role for IFN-λ in suppressing the pathogenic activity of neutrophils that are recruited into the skin during HSV-1 infection.

While our studies first focused on the protective effects of IFN-λ signaling in leukocytes, specifically neutrophils, we found that IFN-λ signaling in keratinocytes also was required to restrict HSV-1 skin disease. To investigate the mechanism by which IFN-λ signaling in keratinocytes modulates skin immunity to control HSV-1 skin disease, we performed bulk RNA-seq analysis in primary mouse keratinocytes, as well as in a human epithelial cell line (A549). Our data from A549 cells supports the standard paradigm that the transcriptional response induced by IFN-λ is less potent than the IFN-β response. However, in primary keratinocytes we found that many ISGs were induced equivalently by IFN-λ and IFN-β. These observations are consistent with other transcriptional profiling experiments in primary cells (Caine et al., 2019; Galani et al., 2017) suggesting that experiments in cell lines may underestimate the potency of the IFN-λ response. A notable difference between the IFN-λ and IFN-β responses in keratinocytes was the chemokine CXCL9, which was only weakly induced by IFN-λ compared to IFN-β, consistent with prior work showing that IFN-λ failed to induce expression of CXCL9 and other chemokines, downstream of a lack of STAT1 homodimer formation and IRF1 induction (Forero et al., 2019). Although we found that the IFN-β response was not impacted by the lack of IFN-λ signaling, the IFN-λ response was diminished in both keratinocytes and A549 cells lacking IFN-αβ signaling, suggesting a possible role for cross-talk between the IFN-λ and IFN-αβ responses.

While we found greater accumulation of neutrophils in skin lesions from *Ifnlr1*^-/-^ mice compared to wild-type mice and showed that neutrophils were necessary for the severe skin pathology observed in *Ifnlr1*^-/-^ mice, the mechanism by which IFN-λ modulates neutrophil function to protect against HSV-1 skin disease remains unclear. We found no difference in CD63 expression in neutrophils from *Ifnlr1*^-/-^ mice compared to wild-type mice, suggesting that IFN-λ does not act to inhibit degranulation in this system. We did find that *Ifnlr1^-/-^* mice exhibited increased expression of the lectin LFA-1, which promotes tissue recruitment and retention of neutrophils, consistent with a model where IFN-λ signaling reduces neutrophil retention in HSV-1 skin lesions. Accordingly, *Cxcl9* was among the genes with reduced induction by IFN-λ compared to IFN-β in primary keratinocytes, CXCL9 levels in skin lesions were higher in the absence of IFN-λ signaling, and depleting CXCL9 ablated the enhanced neutrophil infiltration and large skin lesion phenotypes of *K14*-Cre-*Ifnlr1*^-/-^ mice. Altogether these data support a model where in the absence of IFN-λ signaling, keratinocytes overproduce CXCL9, stimulating excessive neutrophil infiltration and retention into HSV-1 infected skin, exacerbating skin disease severity.

In addition to HSV-1, we also found a protective role for IFN-λ in HSV-2 skin infection. Like HSV-1, HSV-2 produced larger skin lesions in the absence of IFN-λ signaling, although the average severity of HSV-2 lesions was greater than HSV-1 and the kinetics of the infection were slower. The fact that we observed a similar role for IFN-λ for both HSV-1 and HSV-2 supports a general role for IFN-λ in controlling viral skin infections; future studies will investigate the protective effects of IFN-λ against other viral pathogens that target the skin, including poxviruses, papillomaviruses, and arboviruses, as well as bacterial, fungal, and protozoan skin pathogens. We also evaluated whether IFN-λ protected against HSV vaginal infection, since the vaginal epithelium is a plausible site for IFN-λ activity and IFN-λ has been reported to protect against vaginal HSV-2 and Zika virus infections (Ank et al., 2008; Caine et al., 2019). HSV-1 vaginal infection typically does not produce overt disease in mice, whereas HSV-2 produces disease signs that can be evaluated in addition to measuring viral loads. Prior work with HSV-2 vaginal infection in wild-type mice showed that neutrophil recruitment to the vagina worsened disease severity (Lebratti et al., 2021), further supporting the premise for a protective role for IFN-λ at this epithelial barrier. However, we found no difference in overt vaginal disease in *Ifnlr1*^-/-^ mice compared to wild-type mice infected with HSV-2. These results are consistent with our previous observations with Zika virus, where we found no difference in vaginal wash viral loads from *Ifnlr1*^-/-^ mice compared to wild-type mice (Lopez et al., 2022). Additionally, our data are consistent with other reports for Zika virus and HSV-2 vaginal infections, as these experiments only found protective effects for IFN-λ signaling when pretreating mice intravaginally with recombinant IFN-λ or Toll-like receptor 3 (TLR-3) or TLR-9 agonists (Ank et al., 2008; Caine et al., 2019), as opposed to evaluating phenotypes in *Ifnlr1*^-/-^ mice. Overall, these observations suggest any protective effect of IFN-λ in the vagina may be context-dependent and likely does not serve to restrict viral replication. The difference in phenotypes for IFN-λ signaling against HSV-2 infection in the skin and the vagina also underscores possible differences in mechanisms of immunity at these different epithelial barriers.

Although our work has focused on IFN-λ-mediated protection from HSV-1 in context of healthy skin, it will be interesting to evaluate these effects in the context of dermatological conditions such as atopic dermatitis (also called eczema). Patients with atopic dermatitis are susceptible to severe HSV-1 skin infections (eczema herpeticum) (Wollenberg et al., 2003). Notably, atopic dermatitis skin lesions exhibit diminished IFN-λ production compared to healthy skin from the same patients (Wolk et al., 2013) and injection of recombinant IFN-λ reduced the severity of eczema herpeticum in mice (Kawakami et al., 2017). Atopic dermatitis patients also are susceptible to severe infections by poxviruses, posing a hazard to individuals who require smallpox vaccination (due to risk of exposure to or laboratory work with variola virus, vaccinia virus, or other orthopoxviruses) (Reed et al., 2012; Wharton et al., 2003). We expect that the mechanisms by which IFN-λ restricts HSV-1 skin disease will be relevant to other skin infections, particularly where pathogen-triggered immune pathology produces skin lesions, such as vaccinia virus and *Leishmania* (Novais et al., 2021; Shmeleva et al., 2022).

We found that *Ifnlr1*^-/-^ mice developed larger HSV-1 skin lesions compared to wild-type mice, even though there was no difference in viral loads in the skin, supporting an immunomodulatory, rather than antiviral, role for IFN-λ in the skin. However, in this system, IFN-λ is induced in the skin in response to HSV-1 infection, meaning that the virus has a head start and can employ its arsenal of strategies to antagonize the host IFN response. When we instead pre-treated mice with topical IFN-λ prior to infection, we found lower viral loads in IFN-λ treated mice compared to PBS-treated mice, indicating that IFN-λ can have an antiviral effect against HSV-1 in the skin. IFN-λ has demonstrated therapeutic utility against a variety of viral infections, including hepatitis C virus, hepatitis B virus, and SARS-CoV-2 in human studies, as it is thought to confer protective antiviral activity without the damaging immune pathology induced by IFN-αβ treatment (Feld et al., 2021; Lazear et al., 2019). The skin is an especially attractive site to leverage the protective effects of IFN-λ, due to its easy accessibility for topical administration. Encouragingly, topical IFN-λ treatment restricted HSV-1 corneal infection in mice (Miner et al., 2020). Future studies will investigate the potential for topical IFN-λ treatment as a therapy for HSV and other skin infections.

Altogether, our data provide the first evidence of a skin-specific role for IFN-λ in controlling a viral infection. We detail a mechanism by which IFN-λ signals through both keratinocytes and leukocytes to limit severe HSV-1 skin disease. We found that IFN-λ signaling suppresses neutrophil recruitment to the skin, in part by suppressing CXCL9, and the excess recruitment of neutrophils to the skin in the absence of IFN-λ signaling is the driver of pathology. This keratinocyte-neutrophil axis via CXCL9 provides new insights into the immunomodulatory effects of IFN-λ at barrier sites, including the skin.

## MATERIALS AND METHODS

### Viruses and Cells

Virus stocks were grown in Vero (African green monkey kidney epithelial) cells. Vero and A549 (ATCC# CCL-185) cells were maintained in Dulbecco’s modified Eagle medium (DMEM) containing 5% heat-inactivated fetal bovine serum (FBS) and L-glutamine at 37°C with 5% CO2. Virus stocks were grown in DMEM containing 2% FBS, L-glutamine, and HEPES at 37°C with 5% CO_2_. HSV-1 strain NS was obtained from Dr. Harvey Friedman (University of Pennsylvania) (Friedman et al., 1981). HSV-2 strain 333 was obtained from Dr. Steven Bachenheimer (UNC). Virus stock titers were determined by focus-forming assay on Vero cells. Viral foci were detected using 1:10,000 dilution of αHSV rabbit antibody (Dako) and 1:50,000 dilution of goat αrabbit HRP conjugated antibody, and TrueBlue peroxidase substrate (KPL). Antibody incubations were performed for at least 1 hour at room temperature. Foci were counted on a CTL Immunospot analyzer.

### Mice

All experiments and husbandry were performed under the approval of the University of North Carolina at Chapel Hill Institutional Animal Care and Use Committee. Experiments used 8-12-week-old male and female mice on a C57BL/6 background. Wild-type mice used in Figure 3 and S1 were purchased from Jackson Labs and then bred in house. *Ifnlr1*^-/-^ mice were generated by two breeding schemes. For *Ifnlr1*^-/-^ mice in Figures 1, 5, 6, and 7 mice were generated by crossing *Ifnlr1*^f/f^ mice with mice expressing Cre recombinase under control of an Actin promoter. These mice were bred as Cre hemizygotes to generate mixed litters in which 50% of mice retained IFN-λ signaling and 50% lacked IFN-λ signaling. For *Ifnlr1*^-/-^ mice in Figure 4 and 8, mice were generated by crossing *Ifnlr1*^f/f^ mice with mice expressing Cre recombinase under control of an CMV promoter. After generation, mice were bred as Cre homozygote knockout by knockouts and experimental *Ifnlr1*^-/-^ mice were genotyped periodically to verify *Ifnlr1* knockout status. *K14-*Cre*-Ifnlr1^-/-^*, *Vav-*Cre*-Ifnlr1^-/-^*, *LysM-*Cre*-Ifnlr1^-/-^*, and *Mrp8-*Cre*-Ifnlr1^-/-^* mice were generated by crossing *Ifnlr1*^f/f^ mice with mice expressing Cre recombinase under control of a *Vav* promoter (Jackson Labs #8610), a *K14* promoter (obtained from Dr. Scott Williams, UNC), a LysM promoter (Jackson Labs #4781, obtained from Dr. Jenny Ting, UNC), or a *Mrp8* promoter (Jackson Labs #21614), respectively. Mice were bred as Cre hemizygotes to generate mixed litters in which 50% of mice retained IFN-λ signaling and 50% lacked IFN-λ signaling in specific cell types. After data analysis, experimental mice were genotyped by PCR on tails for both Cre and *Ifnlr1* status*. Ifnar1*^-/-^ mice were obtained from Dr. Jason Whitmire (UNC) then bred in-house. *Ifnar1*^-/-^ *Ifnlr1*^-/-^ DKO mice were generated by crossing *CMV*-Cre *Ifnlr1*^-/-^ and *Ifnar1*^-/-^ mice and bred in house. *Ifnl2/3^-/-^* mice were obtained from Dr. Megan Baldridge (Washington University in St. Louis) and then bred in house.

*Ifnk^-/-^* mice were generated in the laboratory of Dr. Michael Diamond (Washington University in St. Louis). Embryonic stem (ES) cells (C57BL/6 background) were obtained from the knockout mouse project (KOMP). The ES cells contained a LacZ reporter-tagged deletion allele with a neomycin selection cassette (Ifnk^tm1(KOMP)Vlcg^). Mice derived from these ES cells were crossed to mice expressing Cre recombinase under a CMV promoter to excise the neomycin selection cassette. The resulting *Ifnk*^-/-^ mice were bred at UNC for use in experiments. Genetic background was confirmed by miniMUGA genotyping array.

### HSV Skin Infections

One day prior to infection, mice were anesthetized by nose-cone isoflurane and depilated manually by plucking the fur from the right flank. One day later, mice were anesthetized by chamber isoflurane for infections. To perform infections, we abraded the skin of anesthetized, depilated mice with ∼10x closely spaced punctures (over ∼5 mm^2^) using a Quintip skin test allergy needle. Immediately after abrasion, we overlaid 10 µL of viral inoculum (virus + 1% FBS in PBS) and allowed the inoculum to dry while mice were anesthetized.

### PCR Genotyping

*Ifnlr1*, *Cre*, *Ifnl2/3,* and *Ifnk* genotypes were determined by PCR on tail samples (unless otherwise indicated) on DNA extracted using the Quantabio supermix and the following previously described primers:

*Ifnlr1* F_1_5-AGGGAAGCCAAGGGGATGGC-3′, R_1_5′-AGTGCCTGCTGAGGACCAGGA-3′, R_2_5′-GGCTCTGGACCTACGCGCTG-3′; *Cre* F_1_5′-CGTACTGACGGTGGGAGAAT-3′, R_1_5′-CCCGGCAAAACAGGTAGTTA-3′; *Ifnl2/3* F_KO_5′-CACACTGTGGACAGGCCAT-3′, R_KO_5′-ACCAGCCTGAGGTCCCTAGT-3′, F_WT_5-TGGAGTCCAGAGCAGCTTTT-3′, R_WT_5′-TCCACCCAGAAGCAAAGAAC-3′; *Ifnk* F_1_5′-CCTGTGTGTCAGGTTTAC-3′, R_1_5′-TGCCCAACTCCAGGTAGAC-3′, R_2_5′-GTCTGTCCTAGCTTCCTCACTG −3′; *K14-Cre* (used for colony establishment) F_1_5′-CACGATACACCTGACTAGCTGGGTG-3′, R_1_5′-CATCACCCACAGGCTAGCGCCAACT-3′; *Vav-Cre* (used for breeders) F_1_5′-AGATGCCAGGACATCAGGAACCTG-3′, R_1_5′-ATCAGCCACACCAGACACAGAGATC-3′.

### Focus-forming Assay

Viral loads were measured by focus-forming assay in which tissues were weighed, homogenized using beads and DMEM with 2% FBS and 1% HEPES and serially diluted in a round bottom 96-well plate. 100 µL of each dilution was then transferred to monolayers of Vero cells (96 well plate with 2×10^4^ cells/well plated 1 day prior) in duplicate and incubated at 37°C for 1 hour. Thereafter, wells were overlaid with 125 µL methylcellulose (1% methylcellulose in 1X MEM+ 2% FBS, L-Glut, P/S, HEPES) and left to incubate at 37 °C for 20h. After 20h, plates were fixed by adding 4% paraformaldehyde (PFA) 100 µL/well for 60 min at room. After fixing, PFA and overlay was flicked off and the plates were washed 3x with PBS+1%Tween using a handheld plate washer. Foci were detected using 1:10,000 dilution of αHSV rabbit antibody (Dako) and 1:50000 dilution of goat αrabbit HRP conjugated antibody, and TrueBlue peroxidase substrate (KPL). Antibody incubations were performed for at least 1 hour at 24°C. Foci were counted on a CTL Immunospot analyzer.

### Viral Genome Quantification by qPCR

Viral genomes were quantified from pooled DRG samples (L2-L5 from the ipsilateral or contralateral sides) that were homogenized in 500 µL of PBS and beads. DNA was extracted from 200 µL of homogenate using the Qiagen DNeasy Blood & Tissue Kit (#69504). HSV-1 DNA was quantified by TaqMan qPCR on a CFX96 Touch real-time PCR detection system (Bio-Rad) against a standard curve generated by extracting DNA from an HSV-1 viral stock. HSV-1 genomes were detected using the following primers targeting the U_L_23 gene: F primer 5′-TTGTCTCCTTCCGTGTTTCAGTT-3′, R primer 5′-GGCTCCATACCGACGATCTG-3′, and probe 5′-FAM-CCATCTCCCGGGCAAACGTGC-MGB-NFQ-3′ (Ma et al., 2014).

### Lesion Area Quantification and Categorization

To measure HSV lesion areas, mice were anesthetized and photographed using an iPhone camera next to a ruler and an identifying card. Thereafter, images were analyzed using ImageJ in which pixels were converted to millimeters using the reference ruler and then lesions were outlined using the freehand tool and calculated area within the freehand designations were reported. Lesions analyzed 6 dpi were categorized using the following system: <5 mm^2^ = Mild, 5-23 mm^2^ = Moderate, and >23 mm^2^ = Severe.

### HSV-2 Vaginal Infections and Disease Categorization

To infect mice with HSV-2 vaginally, experimental mice were treated 5 days prior with 2 mg DepoProvera (UNC Pharmacy) via subcutaneous inoculation. On the day of infection, mice were inoculated with 10^3^ FFU/10µL of HSV-2 strain 333 intravaginally. Mice were monitored for disease signs starting 4 dpi and categorized using the following scoring system (Lebratti et al., 2021): 0: no inflammation, 1: mild redness and swelling, 2: visible ulceration and fur loss, 3: severe ulceration and signs of sickness, 4: hindlimb paralysis, 5: moribund/dead.

### Bone Marrow Chimeras

Bone marrow chimera mice were generated by lethally irradiating wild-type and *Ifnl2/3^-/-^* mice (2×600 CGy) and injecting 10^7^ bone marrow cells from complementary and reciprocal donors. After irradiation, mice were given sulfamethoxazole (6 mg/ml) in their drinking water for the duration of their life. Mice were infected with HSV-1 and photographed and harvested 6 dpi. Efficient bone marrow transfer was validated at the end of the experiment by PCR genotyping tail, ear skin, and whole blood from each mouse for *Ifnl2/3*.

### Flow Cytometry

For analysis by flow cytometry, tissues were harvested into cold DMEM + 10% FBS and processed using established methods for digestion of whole mouse back skin (Lou et al., 2020). In brief, samples were cut into thirds and incubated overnight at 4°C in 5 mg/mL Dispase II (Sigma) in Hanks Buffered Salt Solution (HBSS). Afterwards, samples were washed 1x with cold PBS and epidermis and dermis were mechanically separated for separate enzymatic digestions. The epidermis was digested in 1 mL 0.05% trypsin (Fisher) for 30 min at 37°C. Separately, the dermis was digested in 3 mL dermal dissociation solution (1 mg/mL Collagenase P (Sigma), 100 µg/mL DNAse I (Fisher) for 1 hour at 37°C. After incubation, matched epidermal and dermal digestions were combined and passed through a 70µm cell strainer and resuspended in DMEM + 10% FBS. Suspensions were then centrifuged at 400xg for 5 min at 4°C and resuspended in PBS + 1% FBS (staining buffer) and stained with fluorescent antibodies.

Fc Block and all fluorescent antibodies were sourced from Biolegend and titrated prior to use. Samples were treated with Fc Block for 10 min at 4°C prior to staining. Afterward, samples were washed 3x in staining buffer and then stained for cell surface antigens for 20 min at 4°C. Cells were then washed 3x, fixed with 2% paraformaldehyde solution and analyzed on an LSRII (BD). Data were analyzed with FlowJo software (v10.8). The following fluorescent antibodies (and specific clones) were used: αCD3 (145-2C11), αEPCAM (G8.8), αCD8a (54-6.7), αCD4 (CK1.5), αCD45 (30-F11), αNK1.1 (PK136), αLy6G (IA8), αCD11c (N418), Fc Block αCD16/32 (93), αCD11b (M1/70), αCD19 (6D5), αLy6C (HK1.4) αCD63 (NVG-2), αLFA-1 (H155-78), Zombie UV Fixable Dye. Gating strategies for neutrophils are shown in Fig. S3. All gating strategies for all cell types and neutrophil phenotyping markers were based upon published immunophenotyping methods (Eichelberger et al., 2019; Lou et al., 2020; Sakamoto et al., 2021).

### Histopathology

Samples were prepared for histology by collecting the entire depilated region of the right flank (lesional and healthy skin), flattening the skin on cardstock, and submerging in 10% Neutral Buffered Formalin for 48 hr at 4°C. After fixation, skins were washed and resuspended in 70% ethanol and stored at 4°C until submission to the UNC Pathology Services Core Facility. To prepare fixed skin for submission, two cuts of skin were taken: 1) a horizontal section through the flank to capture lesion and healthy skin and 2) a vertical cut to capture the diseased dermatome region of skin. Thereafter, samples were stored in 70% ethanol and processed by the core, stained for H&E and HSV-1 (1:1000, Dako), and analyzed by a veterinary pathologist (Dr. Hannah Atkins), who selected representative images and identified regions of interest.

### Neutrophil and CXCL9 Depletions

To deplete neutrophils, mice were injected intraperitoneally 0, 2, and 4 dpi with 250µg in 200µL of αLy6G antibody (Clone 1A8 IgG2a, BioXCell) or with isotype control antibody (BioXCell) or PBS. Systemic depletion of neutrophils was validated at the end of the experiment in each cohort of mice using the gating strategy outlined in supplementary Fig. 3B on splenocytes. To neutralize CXCL9, mice were injected intraperitoneally 0, 2, 4 dpi with 300 µg in 200 µL of αCXCL9 antibody (Clone MIG-2F5.5 IgG, BioXCell).

### CXCL9 ELISA

To measure the concentration of CXCL9 in the skin, lesional and healthy skin were collected from wild-type and *Ifnlr1^-/-^* mice 6 dpi. Samples were weighed and then homogenized using beads and resuspended in 1 mL of DMEM. CXCL9 was measured from homogenized skin supernatants using the MIG/CXCL9 Mouse ELISA Kit (ThermoFisher, #EMCXCL9) using optical density read at 450nm and fit against the standard curve according to manufacturer protocol.

### IFN-λ Topical Pre-treatment

To prophylactically treat mice with IFN-λ, mice were depilated by plucking 2 days prior to infection. The next day, 5 µg of recombinant murine IFN-λ3 (PBL #12820) in 10 µL PBS (or PBS alone) was applied topically to a 5mm^2^ region of skin and allowed to air dry. The treated area of skin was circled with a Sharpie and 24 hours later mice were infected with HSV-1 at the treated site.

### Generating IFN Receptor Knockout A549 Cell Lines

IFN receptor knockout cell lines were generated in an A549 background using CRISPR/Cas9 gene editing. A single guide RNA targeting exon 2 of IFNAR1 (5’-CACCGTAGATGACAACTTTATCCTG-3’), exon 3 of IFNLR1 (5’-CACCGACAAGTTCAAGGGACGCGTG-3’), and a non-targeting control (5’-CACCGGTATAATACACCGCGCTAC-3’) were cloned into the lentiCRISPRv2 vector (Addgene). The gRNA construct was then co-transfected with two packaging plasmids (pMD2.G and pSPAX2, Addgene) into HEK293T cells to produce lentiviruses. A549 cells were transduced and selected using 2.5 µg/ml puromycin for two weeks. Surviving cells were then single cell sorted into 96 well plates using a FACS ARIA II flow cytometer. Single clones of knockout monoclonal cultures were confirmed via Sanger sequencing and used for subsequent experiments.

### Primary Mouse Keratinocyte Isolation

Keratinocytes were isolated from adult male and female mice of indicated genotypes using published protocols (Li et al., 2017). In brief, mice were euthanized and whole tails were harvested into cold PBS. Tail skin was peeled off, cut into thirds, and digested overnight at 4°C in 4 mg/mL Dispase II (Sigma) while rotating. The epidermis was then mechanically peeled from the dermis and placed basal side down into 0.05% trypsin to shake for 20 min at room temperature. Afterwards, 7 mL of DMEM with 10% FBS was added and the epidermis was rubbed basal side down into the bottom of a petri dish to release basal keratinocytes. Cell suspensions were collected, passed through a 70 µm cell strainer, and pelleted at 400xg for 5 min. Cell pellets were resuspended in DermaLife K Keratinocyte Medium without antibiotics (Lifeline, Cat #LL-007) to 1×10^5^ cells/mL and seeded 1mL/well in 12-well plates. One adult mouse tail generally yields ∼1×10^5^ cells. Keratinocytes were grown to confluence, ∼3-4 days, with media changes every other day, prior to experimental use.

### IFN treatments in Cell Culture

Primary mouse keratinocytes in 12-well plates were grown to confluence for 3-4 days after isolation. Recombinant murine IFN-β (5 ng/mL; PBL #12401) or IFN-λ3 (5 ng/mL; PBL# 12820) was resuspended in fresh keratinocyte growth media and 1 mL of IFN suspension or media alone was added to cells. 24 hours later, supernatant was removed, 350 µL of Buffer RLT (Qiagen) was added to each well, and samples were stored at −80 °C until RNA extraction. A549 cells were treated similarly at confluence in 6-well plates with recombinant human IFN-β (5 ng/mL; PBL #11415) or IFN-λ2 (50 ng/mL; PBL #11720) or media only, resuspended in A549 growth media. Then, 8 hours later, supernatant was removed, 350 µL of Buffer RLT (Qiagen) was added to each well, and samples were stored at −80°C until RNA extraction.

### RNA extractions and qRT-PCR

RNA from primary keratinocytes and A549 cells was extracted with the Qiagen RNAeasy minikit. Predesigned primer/probe sets (Integrated DNA Technologies) were used to detect murine *Ifit1* (Mm.PT.58.32674307) and *ActB* (Mm.PT.39a.22214843.g) and human *IFIT1* (Hs.PT.56a.20769090.g) and *ACTB* (Hs.PT.39a.22214847). Transcripts were measured by TaqMan one-step qRT-PCR on a CFX96 Touch real-time PCR detection system (Bio-Rad) and were reported as relative differences in threshold cycle (Ct) values between ISG and actin as a housekeeping gene using the 2^-ΔΔCt^ with reference to the average of media-treated, wild-type primary keratinocytes or RenLuc A549s for each sample.

### RNA Sequencing

RNA extracted from A549 cells and primary keratinocytes with the indicated IFN treatments was prepared for RNA sequencing by the UNC High Throughput Sequencing Facility using the Kapa mRNA stranded library prep kit. Primary keratinocytes were sequenced using NovaSeq6000S4 XP Paired End 2×100 and A549 cells were sequenced using NovaSeq6000S4 XP Paired End 2×150. Sequences were analyzed using CLC Genomics Workbench version 23.0.4 (Qiagen) with references to *Mus musculus* reference genome sequence and annotation mm10 (Ensembl GRCm39.110) and *Homo sapiens* reference genome sequence and annotation hg38 (Ensembl GRCh38. 110) for sequences from primary keratinocytes and A549 cells, respectively, using default settings. Gene lists were then analyzed using principal component analysis and volcano plot tools within CLC Genomics Workbench. Differentially expressed genes were defined by having a false discovery rate (FDR) < 0.05 and an absolute value Log_2_ Fold Change > 1. Statistics were calculated within CLC Genomics Workbench and volcano plots were prepared for visualization using Graphpad Prism 9 and principal component analysis plots using CLC Genomics Workbench. Raw and processed RNA sequencing data is available under the GEO superseries accession number GSE242171.

### Statistics

All statistics (apart from RNAseq data) were calculated using GraphPad Prism 9 or XLSTAT 2022 Plugin. Lesion area, viral loads, and vaginal disease score were analyzed using a Mann-Whitney U test and adjusting for multiple comparisons using the Holm-Šídák method when appropriate. Flow cytometry was analyzed using an unpaired t-test. HSV-1 skin disease severity was analyzed using the Cochrane-Armitage Method and adjusting for multiple-comparisons using the Benjamini and Hochberg method when appropriate. Survival curves were analyzed using the Mantel-Cox Log Rank test.

## ACKNOWLEDGEMENTS

This work was supported by R01 AI139512 and R01 AI175708 (H.M.L.). D.T.P. was supported by T32 AI007419 and F31 AI167502. IM and JKW were supported by R01 AI138337, R01 AI131685, R01 AI143894 to JKW, and R21 AI163606 to IM, and a DoD award W81XWH2110919 (JKW). The UNC flow cytometry core facility is supported by a NCI Center Core Support Grant (5P30CA016086). We gratefully acknowledge the technical support from the UNC High Throughput Sequencing Facility. This facility is supported by the University Cancer Research Fund, Comprehensive Cancer Center Core Support grant (P30-CA016086) and the UNC Center for Mental Health and Susceptibility grant (P30-ES010126). The authors have no competing financial interests.

## SUPPLEMENTARY MATERIALS

Supplementary Figures 1-4

Supplementary Table 1: RNAseq data from A549 cells

Supplementary Table 2: RNAseq data from primary keratinocytes

---

**Supplementary Figure 1.**
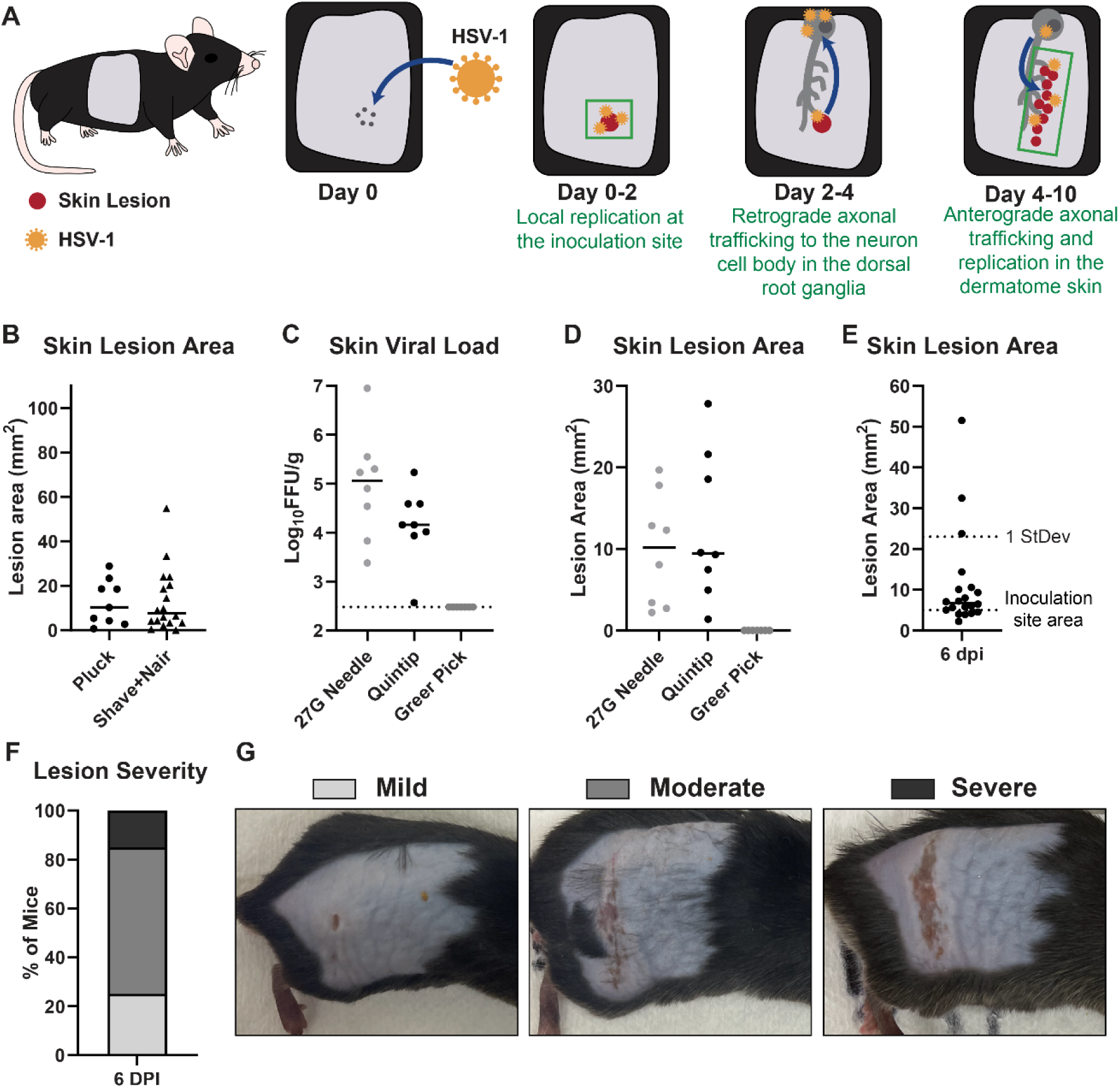
Optimizing an HSV-1 skin infection model. **A.** HSV-1 skin infection model. **B-G.** 8-12 week-old male and female wild-type (C57BL/6) mice were infected with 10^6^ FFU of HSV-1 strain NS and dermatome lesions were analyzed 6 dpi. **B.** Mice were depilated by manual plucking or by shaving plus Nair 1 day prior to infection. **C-D.** Depilated right flank skin of mice was abraded prior to inoculation using 40 scratches with a 27G needle, 10 punctures with a Quintip allergy needle, or 1 puncture with a multi-pronged Greer pick allergy needle. **C.** Skin lesion viral loads were measured by FFA. **D.** Skin lesions were photographed and areas measured using ImageJ. **E-G.** 20 mice were depilated by manual plucking and infected 1 day later using a Quintip allergy needle. **E.** Skin lesions were photographed 6 dpi and lesion areas measured using ImageJ. The standard deviation of lesion areas was 12 mm^2^. **F.** Lesion severity was categorized based on lesion area: <5 mm^2^ = Mild, 5-23 mm^2^ = Moderate, >23 mm^2^ = Severe. **G.** Representative images of each skin disease severity category.

**Supplementary Figure 2.**
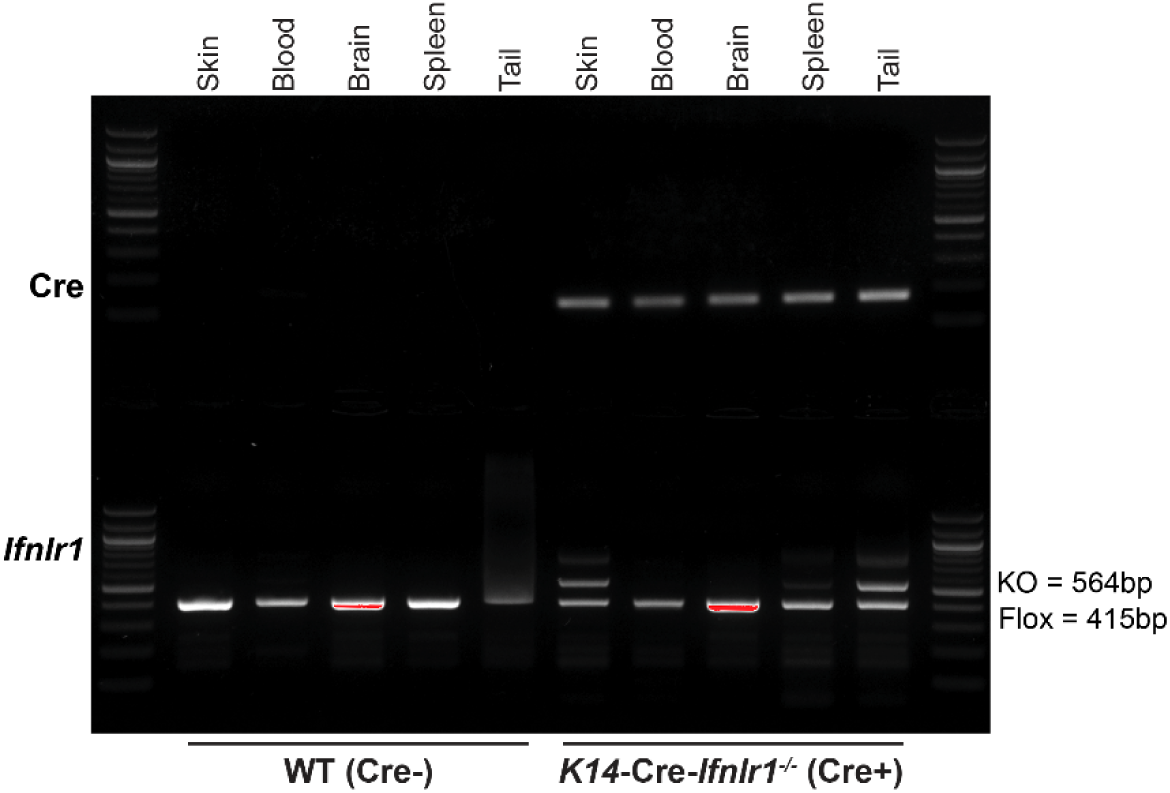
Validation of keratinocyte-specific *Ifnlr1* conditional knockouts. Tissues (flank skin, blood, brain, spleen, tail) were harvested from 8-week-old *K14*-Cre- and *K14*-Cre+ littermates, both carrying homozygous floxed alleles of *Ifnlr1*. Cre and *Ifnlr1* genotyping was performed by PCR.

**Supplementary Figure 3.**
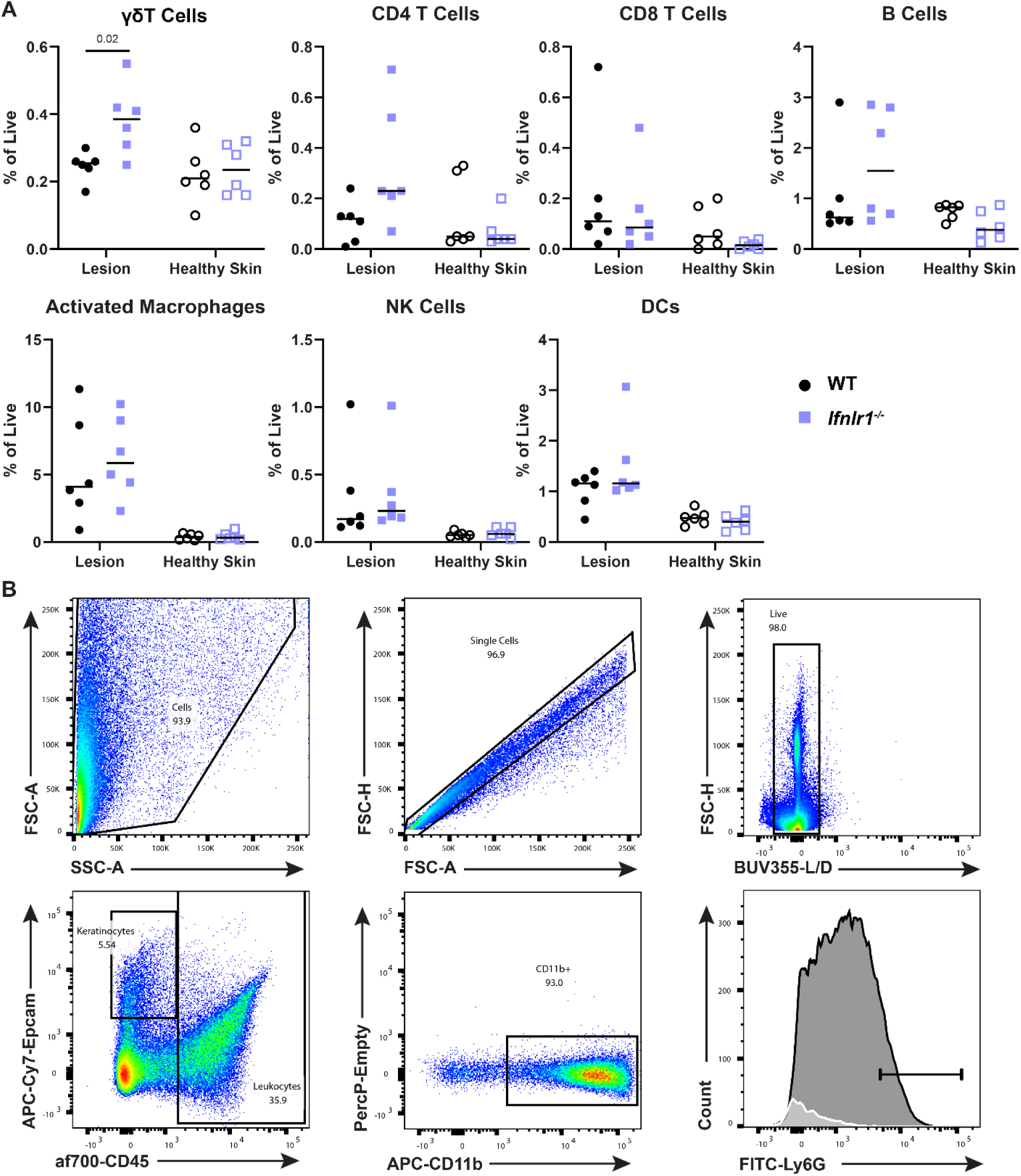
Immunophenotyping of skin tissues from WT and *Ifnlr1^-/-^* mice. **A.** 8-12-week-old male and female WT and *Ifnlr1^-/-^* mice were infected with 10^6^ FFU of HSV-1. Dermatome skin lesions and adjacent healthy skin were collected 6 dpi, processed to single cell suspensions, and analyzed by flow cytometry. All cell populations were normalized to the total live cell population in each sample (Zombie UV-). All leukocytes were CD45+ and the frequencies of leukocyte populations were determined as follows: γδ T cells (CD3+, γδTCR+), CD4 T cells (CD3+, CD4+), CD8 T cells (CD3+, CD8+), B Cells (CD3-, CD19+), dendritic cells (CD11c+; DC), activated macrophages (CD11b+, Ly6G-, Ly6C high), and NK cells (NK1.1+). Differences in leukocyte populations were compared by unpaired t test, with *P* < 0.05 considered to be significant. These mice are the same as those in Fig. 6. **B.** Representative gating strategy to define neutrophil populations.

**Supplementary Figure 4.**
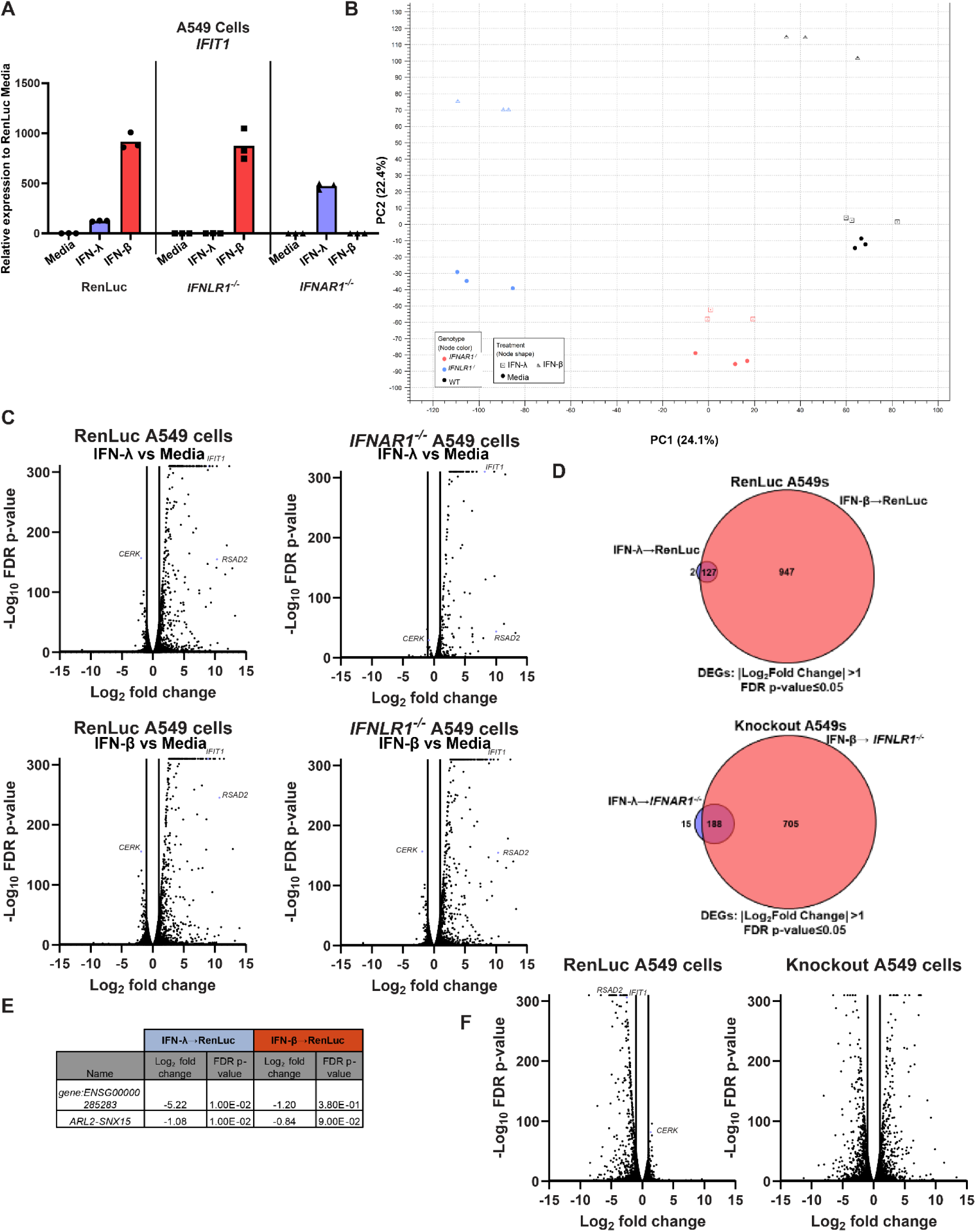
IFN-λ and IFN-β induced transcriptional responses in A549 cells. *IFNLR1*^-/-^, *IFNAR1*^-/-^, and control (renilla luciferase) A549 cells were treated with IFN-λ2 (50 ng/mL), IFN-β (5 ng/mL), or media for 8 hours. **A.** RNA was extracted and IFN-stimulated gene expression was measured by qRT-PCR. *IFIT1* expression is shown relative to *ACTB* (housekeeping gene) and normalized to expression in media-treated Ren-Luc cells. Results represent 3 samples from one experiment. **B-F.** RNA was analyzed by RNAseq (NovaSeq6000S4 XP Paired End 2×100). **B.** Principal component analysis for all analyzed samples. **C.** Volcano plots showing differentially expressed genes after IFN-λ or IFN-β treatment, compared to media-only treated cells. Differentially expressed genes were defined as having a |Log_2_Fold Change| >1 and an FDR p-value ≤ 0.05. **D.** Venn diagram showing DEGs induced by IFN-λ and IFN-β in control or knockout cells. **E.** Table of IFN-λ-specific DEGs, their Log_2_Fold Change, and FDR p-value for IFN-λ and IFN-β-treated control cells. **F.** Volcano plots showing DEGs after IFN-λ treatment compared to IFN-β treatment in control and receptor knockout cells.

